# Information processing in the Hand Laterality Judgement Task: Fundamental differences between dorsal and palmar views revealed by a “Forced Response” paradigm

**DOI:** 10.1101/2025.03.17.643645

**Authors:** Marcos Moreno-Verdú, Baptiste M Waltzing, Siobhán M McAteer, Elise E Van Caenegem, Gautier Hamoline, Robert M Hardwick

**Author notes:** Correspondence: Dr. Marcos Moreno-Verdú.

## Abstract

Imagining performing movements (motor imagery) has broad applications from fundamental neuroscience to sports and rehabilitation. However, measuring motor imagery ability is challenging due to its covert nature. While the Hand Laterality Judgement Task (HLJT) has been investigated as a measure of implicit motor imagery ability, our understanding of mechanisms underlying performance of the task is limited. We used a ‘forced response’ paradigm to study the time-course of information processing in the HLJT. Participants (N=54) performed a modified HLJT where the time they had to process the stimulus was manipulated on a trial-by-trial basis, allowing us to reconstruct the time-course of information processing. Generalised Additive Mixed Models assessed the relationship between processing time and accuracy, which varied across rotation angles (0° to 180° in 45° steps), hand views (dorsal or palmar) or directions (medial or lateral). Stimulus rotation substantively increased the time needed to produce a correct response, although this effect was non-monotonic. Computational modelling confirmed a crucial interaction between hand view and rotation angle, identifying fundamental differences in processing for palmar stimuli with more extreme rotations (≥135°) compared to other stimuli. Finally, a ‘biomechanical constraints’ effect (i.e. faster processing of medial vs laterally rotated stimuli) was present in both views, but was only statistically significant in palmar views, again suggesting differences in processing palmar and dorsal stimuli. These results improve our understanding of the cognitive processes underlying the HLJT and may have broader importance for our understanding of mental processes implicated in motor imagery.

## INTRODUCTION

Motor imagery is the mental simulation of movement in the absence of physical execution (Jeannerod, 2001). Motor imagery spans across various disciplines, from fundamental neuroscience to sports science and rehabilitation (Guerra et al., 2017; Guillot et al., 2021; Ladda et al., 2021; Paravlic et al., 2018). The ability to perform motor imagery can vary between individuals (Floridou et al., 2022; Guillot et al., 2008), but this ability is challenging to measure because of the covert nature of multiple cognitive processes occurring during motor imagery (Cumming & Eaves, 2018).

The Hand Laterality Judgement Task (HLJT) has been proposed as a tool for measuring imagery ability (Heremans et al., 2013; McAvinue & Robertson, 2008) and has also been considered an “implicit” measure of mental representation of the body in action, as opposed to paradigms that require greater action monitoring (Brusa et al., 2023; Scarpina et al., 2019). In this task, the individual has to determine whether a picture of a rotated hand corresponds to the right or left side (Parsons, 1987). The task has been used as the paradigmatic example to argue that motor imagery is implicitly used in body recognition tasks. This model posits that the individual will use the mental representation of their own hand to decide the laterality (Parsons, 1987; Sekiyama, 1982).

Behavioural evidence has repeatedly found a phenomenon which differentiates the HLJT from mental rotation of other objects, namely the ‘biomechanical constraints’ effect (Parsons, 1994). The effect illustrates how anatomical restrictions can influence the mental rotation process: when the hand image is rotated in directions that are more easily physically achieved (i.e., towards the midline or medial direction), the time needed to process the stimulus decreases, compared to images rotated towards anatomically awkward angles (i.e., away from the midline or lateral direction). This phenomenon has been consistently replicated across differing HLJT paradigms (Bek et al., 2022; Conson et al., 2021; Ionta et al., 2007; Meng et al., 2016), and has not been found in mental rotation tasks using letters or alphanumerical symbols as stimuli (Bek et al., 2022; ter Horst et al., 2012). These findings have been used to claim that the biomechanical constraints effect represents a hallmark of the use of motor imagery to complete the task (Bek et al., 2022; Hoyek et al., 2014; Meng et al., 2016; Vannuscorps & Caramazza, 2016).

Critical knowledge gaps are still present regarding how stimuli are processed during the HLJT. For example, previous research has shown an interaction between the degree of rotation and the view of the hand that is presented, yet the nature of this interaction is not well understood. There is behavioural (Conson et al., 2020, 2021; Hoyek et al., 2014) and neurophysiological (Meng et al., 2016; Zapparoli et al., 2014) work suggesting that hand images in a palmar view (i.e., the palm facing up) are processed in different ways than hand images in a dorsal view (i.e., the dorsum facing up). Some authors (Brady et al., 2011; Nagashima et al., 2019) have suggested that the difference may lie in the fact that they might elicit different processing strategies, where processing based on spatial/visual aspects largely occurs for dorsal views (i.e., using an allocentric/third-person reference frame) and processing based on anatomical/motor aspects occurs for palmar views (i.e., using an egocentric/first-person reference frame). In fact, previous work has shown that when hand images display an orientation more plausible from a third-person perspective than from a first-person perspective, as in a rotation angle of 180° in the dorsal view, a mirror-like mapping is automatically generated in the brain (Shmuelof & Zohary, 2008). In other words, dorsal views might more easily trigger an allocentric strategy when stimuli are rotated to a greater extent and closer to a mirror-like position, whereas palmar views would more easily trigger an egocentric strategy regardless of the rotational angle (Bek et al., 2022). However, there are no studies that have formally investigated this hypothesis in the HLJT. Here, using a combination of ‘forced response’ behavioral measurements and computational modelling, we provide the first empirical evidence to support this proposal. Our results indicate that palmar hand stimuli presented at more extreme rotations are processed in a fundamentally different manner from other hand stimuli, which we attribute to a differences between separable allocentric and egocentric modes of processing.

The possibility of differing processing strategies in the palmar and the dorsal view is supported by compelling evidence suggesting that the strength (or even the presence) of the ‘biomechanical constraints’ effect can depend largely on the view of the hand that is presented (Bek et al., 2022; Conson et al., 2021; Hoyek et al., 2014; Mibu et al., 2020; Nagashima et al., 2019). Indeed, hand images in the palmar view typically trigger this effect, whereas hand images in the dorsal view typically show a much weaker effect or do not produce it at all. This could be indirect evidence of motor-based processing occurring only for the palm of the hand (Conson et al., 2021). Understanding the underlying cognitive subprocesses of this medial-to-lateral advantage is useful to determine which specific stimuli are more likely to elicit the different processing strategies, a knowledge that would be more complete if the information processing time-course is studied as a whole, instead of only studying its end-point through classical reaction time paradigm. In our study, we approached this experimentally by modifying the traditional HLJT into a ‘forced response’ protocol, which allowed us to reconstruct the time-course of the ‘biomechanical constraints’ effect. This information is crucial to use the HLJT as a potential motor imagery ability index and increase our understanding of the possible use of motor imagery in this task, which in turn will have an impact on its use in applied contexts.

As indicated above, the traditional HLJT paradigm effectively considers two separate but inherently linked dependent variables (i.e. reaction time and accuracy), and therefore it has a limited ability to study how the stimulus is processed, as speed-accuracy trade-offs can interfere with the dynamics of information processing (Wickelgren, 1977). Moreover, simple reaction time paradigms, although straightforward, only provide data from the endpoint of the processing time-course. Several methods have been proposed to overcome these limitations, both experimentally (Katsimpokis et al., 2020) and statistically (Davidson & Martin, 2013), but other behavioural paradigms could also be useful to provide insight into the time-course of processing during this task. In this scenario, ‘forced response’ paradigms, which ‘force’ individuals to respond after a specific amount of time (i.e. effectively manipulating the duration of stimulus presentation as an independent variable) could help to unveil behavioural signatures typically observed in the HLJT. Forced response paradigms provide a valuable means to reconstruct the time-course of information processing during a task, assessing accuracy as a function of the time allowed to process the stimulus. The forced response approach therefore allows researchers to study the time-course of behaviour during stimulus processing with fine-grained precision (Haith et al., 2016). As an example, forced response protocols enable researchers to understand the evolution of ‘basic’ effects in the HLJT, like the effect of stimulus rotation, which is still not fully understood. Many studies in the literature from general mental rotation paradigms show this effect linearly increases reaction time (i.e., there is a monotonic and proportional relationship between rotation angle and reaction time), and is the main driver of information processing in these tasks. Nonetheless, the HLJT is a particular paradigm where differences in the effect of rotation angle have been reported. For instance, previous studies have found non-linear increases in reaction times for this task (which in some cases are reflective of the ‘biomechanical constraints’ effect, but in other cases are not) (Bek et al., 2022; Conson et al., 2020; Ionta et al., 2007). However, it is challenging with traditional reaction time paradigms to ascertain whether this particular difference is due to actual differences in processing, or artifacts caused by simple shifts in the speed-accuracy trade-off (i.e. a participant may respond more slowly to maintain accuracy, or respond faster at the cost of making more errors).

This paper studied the time-course of information processing during the HLJT. We used a ‘forced response’ paradigm to analyse how factors such as stimulus rotation or hand view affect performance during the task, how they interact with each other, and their influence on the ‘biomechanical constraints’ effect. We hypothesised that information processing is primarily driven by stimulus rotation, and that a strong interaction between rotational angle and hand view would be evidenced. Additionally, we predicted that a medial-to-lateral advantage (consistent with a ‘biomechanical constraints effect’) would only be evident for palmar views, as they are more likely to produce an egocentric (i.e., motor-based) strategy. Using computational modelling, we further characterised different response profiles for hand views and rotation angles, which is key to understand how information processing strategies might be used in this task. Overall, our study stepped beyond current limitations of behavioural paradigms to increase our understanding of information processing in the HLJT.

## METHODS

### Participants

Healthy individuals aged 18-35 years, with normal or corrected-to-normal vision and no history of neurological damage participated. All participants self-described as right-handed. Their self-reported motor imagery ability was assessed using an electronic adaptation of the Movement Imagery Questionnaire-3 (MIQ-3 – see below for further details) (Williams et al., 2012). Participants were recruited via the crowdsourcing platform Prolific (https://www.prolific.com/), and the study was run completely online. Initially, 106 participants were enrolled to participate, of which 73 completed the study. After applying data quality exclusion criteria (see Data Curation for details), data from 54 participants were analysed.

Participants were randomly allocated to one of two experimental groups. The “Dorsal Group” and “Palmar Group” responded to separate sets of stimuli presenting the hand in either a dorsal or palmar view, respectively. Their characteristics are summarised in Table 1.

**Table 1.** Participants’ characteristics according to the experimental group.

| Characteristic | Dorsal Group<br>(N = 28) | Palmar Group<br>(N = 26) | Difference between<br>groups |
| --- | --- | --- | --- |
| Age<sup>1</sup>, mean ± SD | 26.14 ± 4.95 | 26.72 ± 5.22 | $t_{(51)} = -0.41, p = 0.682$ |
| MIQ-3, mean ± SD | | | |
| Total score<sup>2</sup> | 58.33 ± 9.34 | 56.15 ± 10.81 | $t_{(51)} = 0.7, p = 0.435$ |
| Kinaesthetic Imagery | 19.18 ± 3.89 | 18.62 ± 4.32 | $t_{(52)} = 0.5, p = 0.616$ |
| Internal Visual Imagery<sup>2</sup> | 18.89 ± 3.71 | 18.81 ± 4.24 | $t_{(51)} = 0.07, p = 0.941$ |
| External Visual Imagery | 20.07 ± 3.53 | 18.73 ± 3.56 | $t_{(52)} = 1.39, p = 0.171$ |
| Sex, n (%) |  |  |  |
| Female | 13 (46%) | 11 (42%) | $\chi^2(1) = 0.001, p = 0.976$ |
| Male | 15 (54%) | 15 (58%) |  |

<sup>1</sup> Data showed from N=25 participants in the Palmar Group due to a technical problem in 1 participant.
<sup>2</sup> Data showed from N=27 participants in the Dorsal Group due to a technical problem in 1 participant.
Note: continuous variables were compared between groups using independent samples t-tests, and categorical variables using $\chi^2$ tests.

All study procedures were approved by the UCLouvain IPSY Ethics Committee (N°: 2024-21). All participants were financially compensated for their time (10€/h).

### General procedure

The overall procedure is shown in Fig. 1A. All participants first completed an electronic version of the MIQ-3 and then proceeded to complete a ‘forced response’ task using a modified version of the HLJT. The experiment was created in PsychoPy2 version 2023.2.3 and run via Pavlovia (https://pavlovia.org/), PsychoPy’s online platform (Peirce et al., 2019). Stimulus size was determined using “PsychoPy units”, where a size of 1 unit corresponds to the height of the screen when viewed in a landscape orientation. Experiment code can be found in the study’s Open Science Framework (OSF) repository (https://osf.io/z6b4d/).

**Figure 1.**
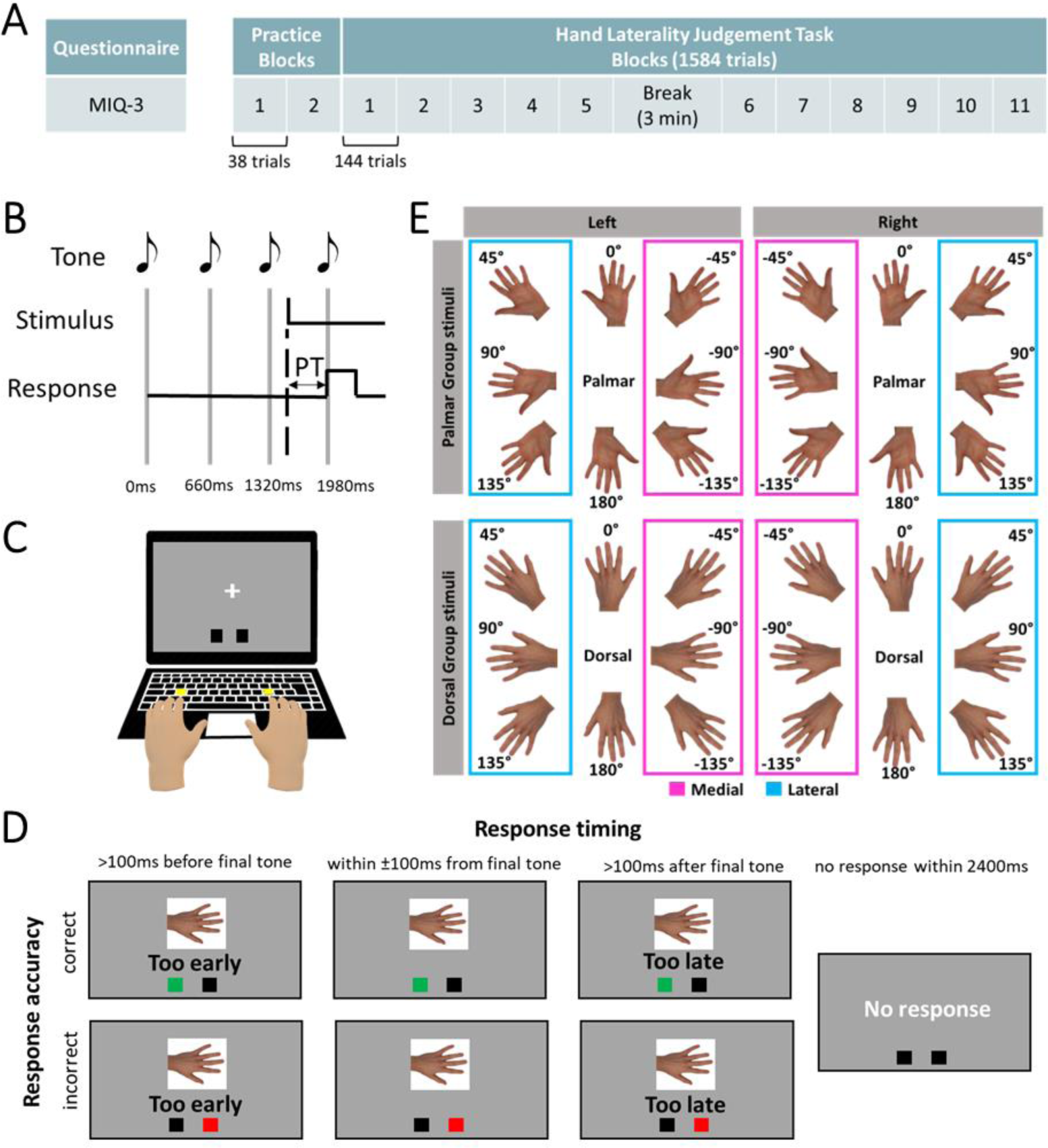
Overview of the procedure of the experiment. Panel. **A** shows the overall structure of the experiment. Participants started by answering the Movement Imagery Questionnaire-3 (MIQ-3), before completing the ‘forced response’ Hand Laterality Judgement Task (including two practice blocks using arrows pointing left or right as stimuli, to familiarise participants with the demands of the forced response paradigm). **Panel B** shows the structure of a ‘forced response’ trial, in which the participant heard 4 tones spaced by 660ms and was asked to synchronise their response with the last tone. The stimulus appeared at a random time between the first and last tone, effectively controlling the amount of “Preparation Time” (PT) the participant had to process the stimulus. **Panel C** shows how participants completed the task; responses were made “bimanually”, using the index fingers to press the ‘S’ and ‘L’ keys for right and left stimuli, respectively. **Panel D** shows examples of possible feedback, which were given on a trial-by-trial basis. Separate feedback was provided based on timing (too early/too late; if the response was in time, no feedback was displayed) and accuracy (box turning green/red for correct/incorrect responses). If no response was provided, a separate screen after the trial ended was provided for 1000ms. **Panel E** shows the experimental stimuli used for the main task (32 unique images, 16 per group). Laterality (right/left) and rotational angle (0° to 180° absolute rotations in increments of ±45°, clockwise or counterclockwise) were used to code medial and lateral directions according to the anatomical position. Angles were encoded such that lateral rotations were positive, and medial rotations were negative.

### MIQ-3

The MIQ-3 was used to quantify motor imagery generation (Williams et al., 2012). The MIQ-3 is a 12-item self-administered questionnaire where participants are asked to imagine four movements in three sensory modalities. The “kinaesthetic” modality focuses on the feelings of the movement, the “internal visual” modality focuses on seeing the movement from a first-person perspective, and the “external visual” modality focuses on seeing the movement from a third-person perspective. For each item the participant is first asked to physically perform the described action, then to imagine it, and finally to rate the ease/difficulty to generate the image on scale from 1 (very difficult to see/feel) to 7 (very easy to see/feel). Scores are then provided as overall (12-84 points) and subscales specific to each sensory modality (4-28 points). An attention check question was interleaved between the actual items, where participants were explicitly instructed to provide a specific response to ensure they were paying attention to the experiment.

### Forced Response Procedure

The main experimental task involved a forced response paradigm (Haith et al., 2016; Schouten & Bekker, 1967; Vleugels et al., 2020; Waltzing et al., 2024). In each trial the time the participant had to respond to the stimulus was systematically manipulated. They heard a series of four tones, each spaced 660ms apart, and were instructed to respond synchronously with the last tone (Fig. 1B). A fixation cross was present in the centre of the screen until the stimulus appeared. The stimulus (dimensions: 0.45 x 0.45 units) was presented in the centre of the screen at a random time between the first and last tone, to effectively manipulate the time the participant had to prepare their response (i.e. their preparation time). Participants made responses bimanually, using the S or L keys to respond using the index finger of the left or right hand, respectively (Fig. 1C).

Participants received separate visual feedback on their response timing and their response accuracy on a trial-by-trial basis. For timing, a threshold of ± 100ms from the last tone was considered correct timing (Fig. 1D). Timing feedback was presented as text below the stimulus (letter height 0.06 units), which showed if the response was too early or too late; if participants were within ± 100ms from the final tone, no text appeared. Accuracy feedback was provided via two small boxes (0.07 x 0.07 units) placed at the bottom of the screen, which turned green if correct and red if incorrect. Note that both types of feedback are critical in this paradigm, as the timing feedback is necessary to get participants to perform the task as instructed, but accuracy feedback is essential to avoid participants focusing only on the timing without trying to respond correctly. Nonetheless, participants were instructed that their highest priority was adhering to the timing constraint, to make sure that the time-course of information processing could be reconstructed appropriately. Adherence to this instruction was used as an exclusion criterion – see below for details. The duration of the feedback message depended on the time of the participant’s response, but each trial had a maximum duration of 2400ms, and therefore this was the maximum feedback duration (if the participant responded just on time, they would have had feedback for approximately 400ms). If after this time the participant had not responded, a message reminding them the need to provide a response was provided for 1000ms.

#### Forced Response Practice Task

Practice blocks were included before the main task to help familiarize participants with the required timing demands. Stimuli in these blocks were images of arrows pointing left or right; participants were instructed to respond with their corresponding hand. The stimuli were presented at 19 possible preparation times (100-1900ms, with steps of 100ms). Participants completed two blocks of 38 practice trials.

#### Forced Response HLJT

In the main task, participants had to judge whether images of hands rotated at different angles depicted a left or right hand. The experimental stimuli were real male hand pictures presenting the hand in a dorsal or palmar view, as shown in Fig. 1E. Images of the left hand were right-hand images that had been mirror reversed. Images were presented at 8 rotational angles in the frontal axis (0°, 45°, 90°, 135°, 180°, −135°, −90°, −45°, positive angles being in the lateral direction and negative in medial direction; Fig. 1E).

For each group, the main task consisted of 16 unique stimuli (8 possible rotations x 2 hands, i.e. left- or right-hand stimuli). We aimed to have a uniform coverage of possible times at which the stimuli appeared. Based on previous studies (Jones et al., 2021), the range decided was 20-1980ms, with steps of 20ms (i.e., stimuli could be presented at preparation times of 20ms, 40ms, 60ms, etc. up to 1980ms). This allowed us to collect data over a broad spectrum of preparation times, from times where participants would have extremely limited time to process the stimuli and should therefore respond at chance level (<300-400ms), and also trials where longer times were available for stimulus processing (> 1500ms). Therefore, stimuli could be presented at 99 possible times (maximum = 1980ms, step = 20ms, hence 1980/20 = 99 times). Overall, each participant completed N = 1,584 trials (16 stimuli x 99 preparation times). The task was divided into 11 blocks (Fig. 1A) to allow for a balanced distribution of preparation times and stimuli across blocks. Each block comprised 144 trials (i.e. each of the 16 unique stimuli were presented at 9 different preparation times). Participants were given the instruction to perform the task in a quiet room and/or wear earphones wherever possible and were given the opportunity to adjust the volume of the computer to their desired level.

Each block had a fixed structure (identical for all participants in both groups) where presentation order was randomized, minimizing the likelihood that two consecutive trials had the same rotational angle (regardless of their laterality), and that no more than 4 consecutive trials presented the same laterality (regardless of their rotational angle). The distribution of preparation times was balanced within and across blocks. The order of blocks was fully randomized for each participant. Between blocks, there was a pause of at least 10 seconds and no maximum limit. To minimise fatigue, there was a mandatory pause of 3 minutes between blocks 5 and 6.

### Exclusion criteria

For each trial, the time at which the stimulus was presented (ms), the time of the participant’s response, and the accuracy of the response were recorded. First, data quality was checked by means of several sequential exclusion criteria. Participants were excluded if they: 1) did not provide a response in at least 75% of trials, as this indicated they did not complete the task as instructed (N=1 in Palmar group; N=0 in Dorsal group); 2) their response time was beyond the limits of ± 100ms from the last tone in more than 50% of trials, excluding participants that did not properly adhere to the instruction to respond synchronously with the last tone (N=4 in Palmar group; N=5 in Dorsal group); 3) their overall accuracy was less than 60%, which indicated not adequately attending to and performing the task (N=5 in palmar group; N=8 in dorsal group). We also had planned to exclude participants if they missed the attention check question of the MIQ-3, but none did. As noted above, this left a final sample of 54 participants (Palmar group: N=26; Dorsal group: N=28).

### Data curation

While the forced response paradigm instructs participants to respond synchronously with the final tone, some natural variability in the exact timing of the response was expected (see Supplementary Figure S1). From the included participants (those meeting all inclusion criteria as described above), we therefore calculated the actual time the participant used to prepare their response (i.e., Preparation Time = response time – stimulus onset time) and used these values in our analyses. This allowed us to analyse all the available trials where a response was provided (even if the response time was beyond the intended limit of ± 100ms), as our analyses considered the actual time participants took to respond, rather than the “prescribed” time imposed by the task. The data was curated to remove trials where implausible Preparation Times were recorded (e.g., Preparation Time being negative, suggesting a response was provided before the stimulus appeared, or Preparation Time being higher than 2400ms, suggesting a technical error occurred given that this was the expected maximum duration of each trial). This represented less than 3% of trials (Supplementary Figure S1).

### Statistical analysis

The analyses were performed in R version 4.3.3 (R Core Team 2024). Data visualisation was performed using the ‘tidyverse’ library and the ‘ggdist’ package (Kay, 2024; Wickham et al., 2019). Scripts and data for analysis are freely available in the study’s OSF repository (https://osf.io/z6b4d/).

Generalised Additive Mixed Models (GAMMs) were used to model the non-linear relationship between Accuracy (binary outcome: correct/incorrect) and Preparation Time (continuous predictor) with by-participant random intercepts (Pedersen et al., 2019; Wood, 2017). The ‘mgcv’ package version 1.9-0 was used to fit the models, with the Restricted Maximum Likelihood (REML) method (Wood, 2011) and low-rank thin plate regression splines (Wood, 2003). All models were run with a binomial distribution for the outcome variable. Details on model building, selection and goodness-of-fit assessment can be found in Supplementary Materials.

First, a model analysing the effects of Rotation Angle and Hand View was built. For this model, we allowed the shape of the Accuracy-Preparation Time relationship to vary across the levels of the absolute Rotation Angle (5 levels: 0°, 45°, 90°, 135° and 180°, regardless of their laterality or direction) and Hand View (2 levels: palmar and dorsal) and their interaction. This was done as we expected strong effects of both rotation and hand view that could vary with time, and could change the form of the speed-accuracy trade-offs. Rotation Angle was treated as an ordinal variable, given we had to discretise it for feasibility purposes, and to keep consistency with previous studies.

Second, a separate model analysing the ‘biomechanical constraints’ effect was built. This model used only those trials where medial and lateral rotations were present (i.e. excluding rotations at 0° and 180°). For this model, we allowed the Accuracy-Preparation Time relationship to vary across the levels of Direction (2 levels: medial and lateral) and Hand View (2 levels: palmar and dorsal) and their interaction, as again we anticipated strong effects of these factors. Therefore, in this model we investigated the effect of Direction by collapsing across all three possible absolute rotation angles (45°, 90° and 135° for lateral and −45°, −90° and −135° for medial; Fig. 1E).

The above models were run collapsing across right and left stimuli, as this was not our main interest. Complementary analyses showed a response preference for right hand stimuli at short Preparation Time (see Supplementary Materials for details).

Unit-level conditional estimates for each of the predictors in the above models were computed and visualised via speed-accuracy trade-offs (i.e. the proportion of correct responses plotted as a function of time used to process the stimulus). These estimates were reported back-transformed from the logit scale to the probability scale. To visualize parameter uncertainty, 95% confidence intervals (95%CI) were obtained by 1,000 bootstrapped replicates. For post-hoc comparisons, the categorical predictors were compared pairwise throughout the range of Preparation Time, while Rotation Angle was compared by pairs and sequentially (i.e. 0° vs 45°, 45° vs 90°, 90° vs 135° and 135° vs 180°, but not 0° vs 90°, etc.). For a more intuitive interpretation, differences in probabilities (via Z-tests) instead of Odds Ratios were computed. P-values were corrected for multiple comparisons using the False Discovery Rate (FDR), fixing the Type I error rate at 0.05.

#### Power analysis

No a priori power analyses were conducted for this study. We aimed to collect data from at least 50 participants (N=25 per group), which is similar to previous studies using Forced Response procedures (Haith et al., 2016; Hardwick et al., 2019; Vleugels et al., 2020). Because we anticipated we would need to reject data from some participants, initially 106 participants were enrolled to participate, of which 73 completed the study and data from 54 participants were analysed (and financially compensated) after checking data quality. Given restrictions on financial resources, we could not collect a larger sample size. After the completion of the study, in order to provide estimates of sample sizes required to replicate the present effects, we performed a prospective, simulation-based power analysis focusing on the interaction between Rotation Angle and Hand View (as it was the most relevant finding of the present study –see Results). The details are explained in Supplementary Materials (Fig. S7-8), which we hope will inform future studies aiming to use Forced Response procedures analysed through GAMMs.

### Computational Modelling

To further characterise the profile of information processing according to Rotation Angle and Hand View, we employed a previously published computational modelling approach (Hardwick et al., 2019). The approach assumes that characteristics of the stimulus are processed after a variable amount of time based on a gaussian distribution (i.e. time T, with variance σ^2^). Responses produced before time T are essentially random (as participants have not had time to process the stimulus and would therefore be guessing the correct answer), while responses after time T are influenced by the presented stimulus.

Simple stimulus-response relationships can therefore be captured using a “single-process” model, whereby the probability of producing a correct response (i.e. the speed-accuracy trade-off; see Fig. 4C) is based on four free parameters and follows a sigmoidal shape with an initial asymptote (reflecting “guesses” with approximately chance-level of accuracy for responses produced before time T), an increase in accuracy (reflecting the cumulative probability of sufficient time to process the stimulus having elapsed, captured by two parameters reflecting the average value of time T and its variance σ^2^), and a final asymptote (representing an approximate plateau at a relatively high level of accuracy).

More complex stimulus-response relationships can be captured using a “dual-process” model (see Fig. 4C). An example is a situation where different features of the stimulus may require differing amount of time to accurately process; for example, in a classic Stroop colour and word task (Stroop, 1935), prepotent, erroneous responses based on the colour of the ink (i.e. Process A) would become available before accurate responses based on the text of the word presented (i.e. Process B). Such relationships can be modelled using two separate processes reflecting the availability of the first available response (i.e. Process A, comprising time T_A_ and its variance σ^2^_A_), and the second available response (i.e. Process B, comprising time T_B_ and its variance σ^2^_B_). The resulting speed-accuracy trade-off could therefore be captured by a model with seven free parameters that captures an initial asymptote (reflecting ‘guesses’ made with approximately chance-level accuracy), an initial decrease in accuracy (reflecting the cumulative probability of erroneous responses based on the initially available Process A, based on the average values of T_A_ and its variance σ^2^_A_), until a plateau point, after which responses based on Process B start to become available, corresponding to an increase in accuracy (reflecting the cumulative probability of correct responses based on Process B, based on the average values of T_B_ and its variance σ^2^_B_), followed by a final asymptote (an approximate plateau at a relatively high level of accuracy).

Separate single-and dual-process models were fit for each level of Rotation Angle across Hand Views. The models were compared via the difference in Akaike’s Information Criterion (ΔAIC), with the lowest AIC indicating evidence in favour of a given model, and a ΔAIC > 10 considered “strong” evidence (Burnham & Anderson, 2004). This analysis was carried out in Matlab version 2023 with a custom-made script.

## RESULTS

### The time-course of information processing followed an expected speed-accuracy trade-off

Overall, 39,960 valid trials were analysed in the Palmar group and 43,243 valid trials in the Dorsal group after data curation. The task took 90 min 21s ± 15 min 22s to complete. See Supplementary Materials for methodological checks.

As a manipulation check, we confirmed that the GAMM identified a non-linear effect of Preparation Time (effective degrees of freedom = 13.87, χ2 = 549.8, p < 2×10^-16^). As expected, the probability of being correct increased with Preparation Time (Fig 2A), which validated the experimental paradigm. Overall (i.e. collapsing across all conditions), accuracy increased over chance level (probability = 0.5) at 413ms, the time at which the lower bound of the 95%CI did not include 0.5 (Fig. 2A; see horizontal line in lower part of plot).

**Figure 2.**
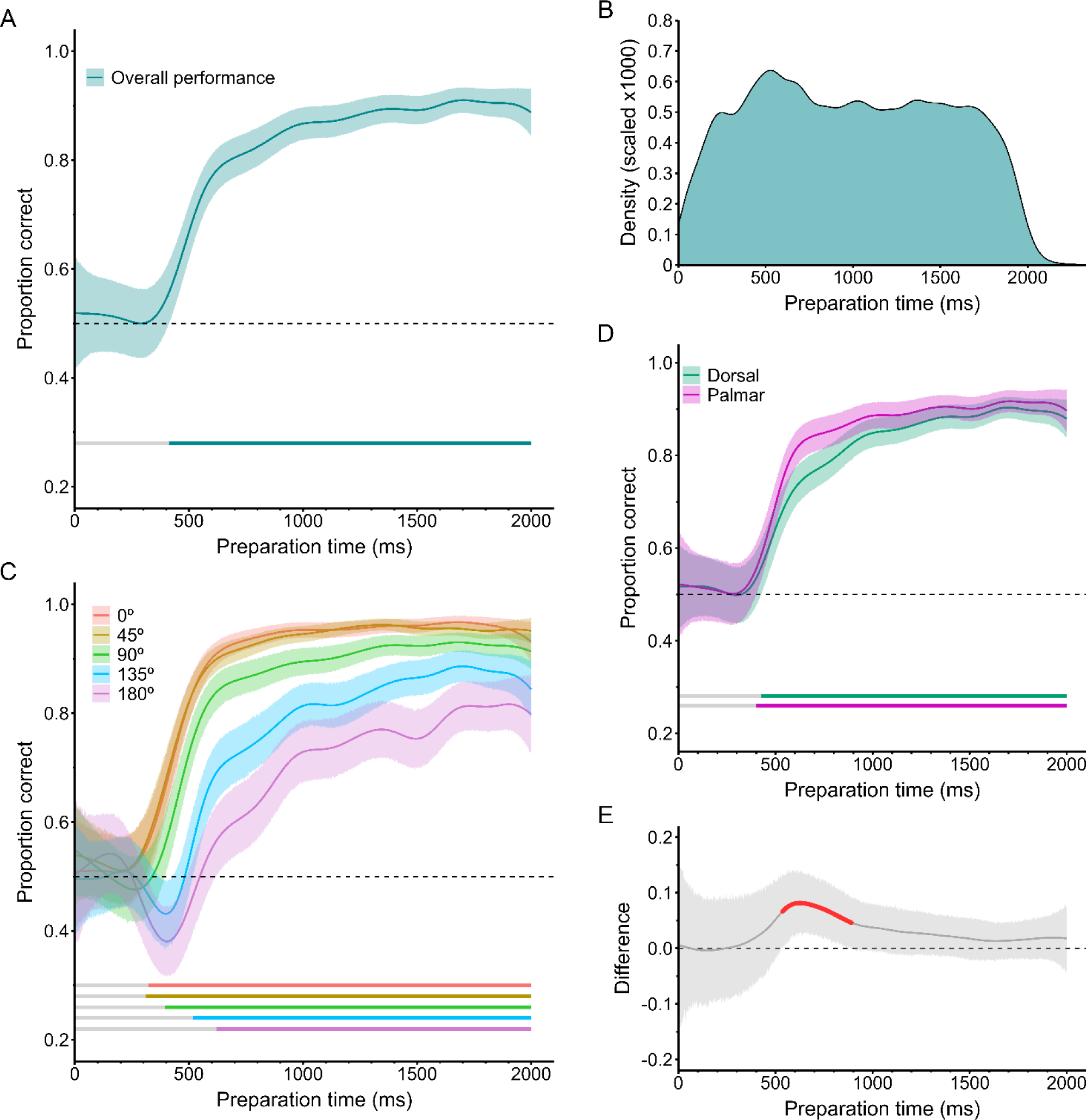
**Panel A** shows the obtained speed-accuracy trade-off collapsing across all conditions. **Panel B** shows a kernel density plot representing the number of trials available for analysis according to the Preparation Time. **Panel C** shows the information processing profile based on Rotation Angle, with all the possible absolute rotations (0° to 180°). **Panel D** shows the information processing profile based on Hand View (palmar or dorsal), and **Panel E** the corresponding difference between Views across Preparation Time. Horizontal lines in the lower part of panels A, C and D indicate the times at which performance in the corresponding condition differed significantly from chance (horizontal dashed line; probability = 0.5). The highlighted red region in Panel E shows where p<0.05, but this did not hold after correction for multiple comparisons (False Discovery Rate method).

### The time needed to process stimuli increased with rotation angle

Overall, as the rotation increased, the time needed to produce a correct response was longer (Fig. 2C). However, throughout the time-course of information processing, the effect of stimulus rotation was non-monotonic (Fig. 3). Post-hoc comparisons did not identify differences in the speed-accuracy trade-off between rotations of 0° and 45° at any time period (difference < 0.01, p_FDR_ > 0.95; Fig. 3A-B). Differences between 45° and 90° were found from 334ms onwards, reaching the peak at 416ms (difference = 0.12 95%CI [0.04, 0.19], p_FDR_ = 0.006) and decreasing afterwards, approaching 0 at the longest Preparation Times, around 1727ms (Fig. 3C-D). Differences between 90° and 135° were found from 371ms onwards, reaching the peak at 484ms (difference = 0.20 [0.12, 0.28], p_FDR_ = 4.14×10^-5^) and linearly decreasing afterwards, with a trend to approach 0 (Fig. 3E-F). Finally, differences between 135° and 180° were found from 524ms to 1162ms and between 1258-1853ms, reaching the peak at 620ms (difference = 0.13 [0.04, 0.21), p_FDR_ = 0.01; Fig. 3G-H).

**Figure 3.**
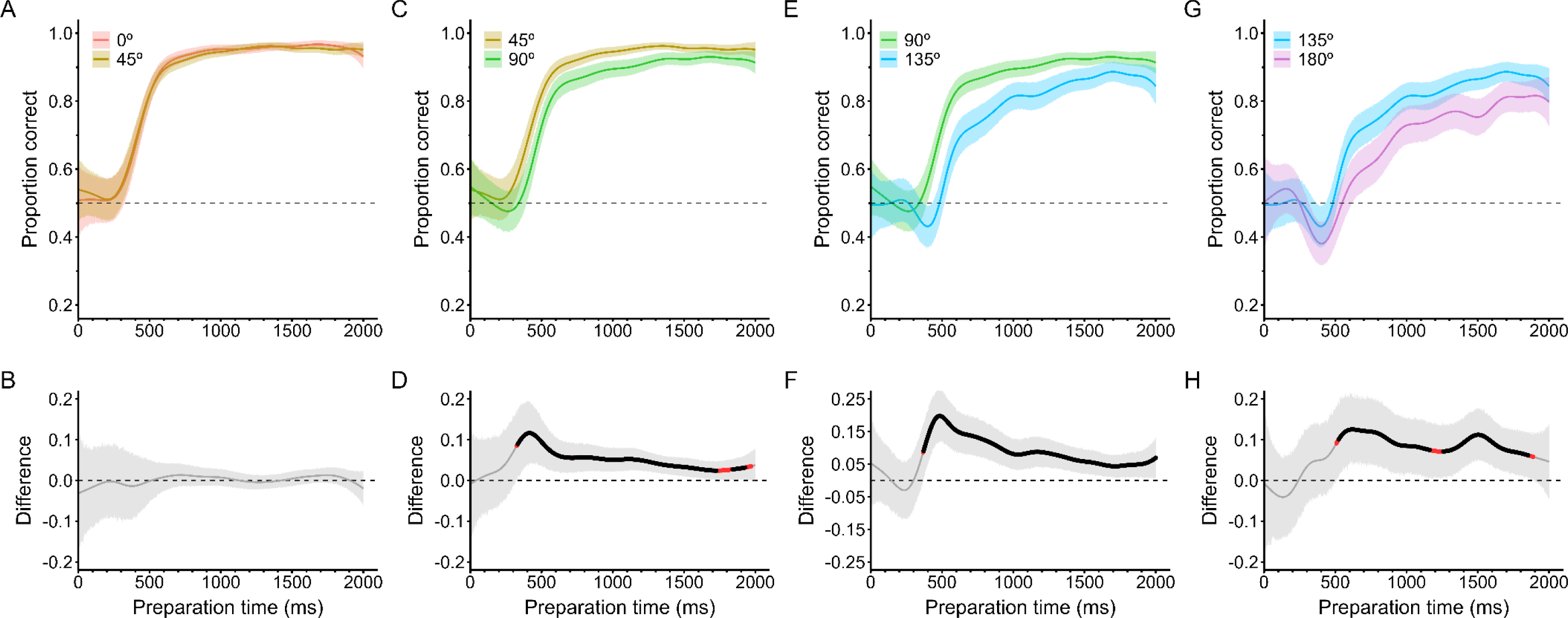
Sequential pairwise comparisons between the absolute Rotation Angles, collapsing across Hand Views. All panels show the information processing profile for each corresponding pair, and below the difference in the probability of being correct between them, across Preparation Time. The difference is computed as Difference = Angle 1 – Angle 2, and therefore positive differences illustrate an advantage of the less rotated stimulus. The highlighted red and black regions in the lower panels show where the difference was p < 0.05 and p_FDR_ < 0.05, respectively.

### Palmar stimuli were processed faster than Dorsal stimuli, although the effect was negligible

Collapsing across all levels of Rotation Angle, the palmar view raised above chance level at 401ms, while this required slightly more time in the dorsal view (426ms; Fig. 2C). Throughout the time-course of information processing (Fig. 2D), the palmar view showed an advantage over the dorsal view in the period 536-891ms, however this difference did not hold after correction for multiple comparisons (p_FDR_ > 0.05). The advantage increased from 536ms onwards, reaching its peak at 626ms (difference = 0.08 [0.03; 0.14], p_FDR_ = 0.089) and reducing afterwards, with the 95%CI gently overlapping 0 from 891ms onwards.

### Greater rotations led to fundamental differences in performance between dorsal and palmar stimuli

Both the computational modelling and GAMM analyses revealed a strong interaction between Rotation Angle and Hand View. This interaction was highlighted by different information processing profiles between views for relatively small rotations compared to more extreme rotations for the palmar but not the dorsal view.

According to the computational model (Table 2), for Palmar stimuli, relatively small rotation angles were best explained by the single-process model (0°, 45°, and 90° all ΔAIC ≤ −12.79), whereas there was strong evidence that more extreme rotations were better explained by the dual-process model (135°: ΔAIC = 39.21 and 180°: ΔAIC = 30.73). By contrast, in the Dorsal view there was evidence in favour of the single-process model for all angles (all ΔAIC ≤ −3.18).

**Table 2.** Summaries of the information processing profiles according to Rotation Angle and Hand View, based on the computational modelling analysis. The difference in Akaike’s Information Criterion (ΔAIC) shows evidence in favour of the single-process model (ΔAIC is negative) or the dual-process model (ΔAIC is positive). Entries in bold highlight where evidence was strongly in favour of the dual-process model (ΔAIC > 10), i.e. palmar views with rotations ≥135°.

| Angle | View | AIC<sub>single</sub> | AIC<sub>dual</sub> | $\Delta AIC$ |
| --- | --- | --- | --- | --- |
| 0° | Palmar | 3258.62 | 3275.1 | -16.47 |
|  | Dorsal | 3903.53 | 3910.95 | -7.42 |
| 45° | Palmar | 6711.62 | 6724.4 | -12.79 |
|  | Dorsal | 7782.72 | 7794.08 | -11.36 |
| 90° | Palmar | 8604.14 | 8617.48 | -13.33 |
|  | Dorsal | 9428.37 | 9434.18 | -5.81 |
| 135° | Palmar | 10776.40 | 10737.22 | **39.21** |
|  | Dorsal | 11990.22 | 12029.11 | -38.89 |
| 180° | Palmar | 5954.89 | 5924.16 | **30.73** |
|  | Dorsal | 7153.15 | 7156.33 | -3.18 |

Results from the GAMM analysis agreed with this finding (Fig. 4). While responses to Palmar stimuli with smaller rotations (0°, 45°, 90°) showed a general increase in accuracy over time, responses to Palmar stimuli with more extreme rotation angles were characterised by an initial dip in accuracy that fell significantly below chance, followed by a later increase in accuracy. For 135°, this occurred in the period 312-465ms (lowest probability = 0.36 [0.3; 0.42], peaking at 398ms), and for 180° it occurred in the period 276-503ms (lowest probability = 0.31 [0.25, 0.37], peaking at 397ms). By contrast, the corresponding responses to Dorsal stimuli never fell below chance level and always showed an increase in accuracy over time (135°: lowest probability = 0.47 [0.38, 0.57]; 180°: lowest probability = 0.45 [0.38, 0.51]). Pairwise comparisons from the GAMM also showed distinct effects of stimulus rotation between hand views (see Supplementary Materials and Figs. S2-S3).

**Figure 4.**
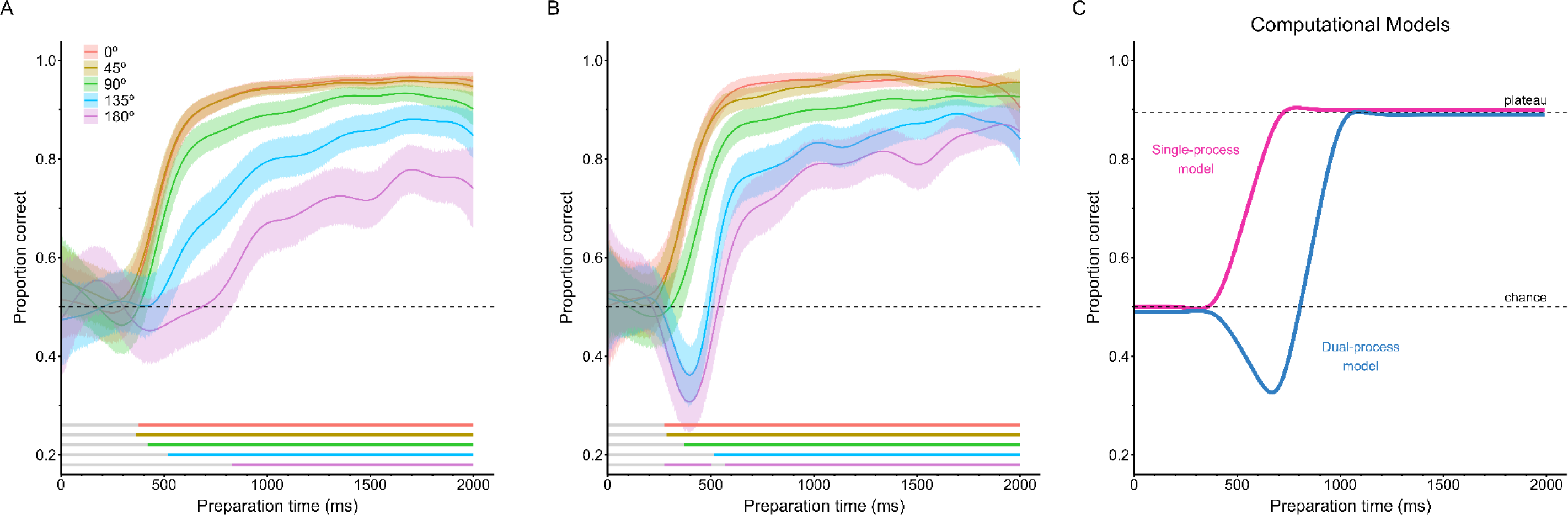
Information processing profiles according to the Rotation Angle for the dorsal view (**Panel A**) and palmar view (Panel B). Horizontal bars show when the condition is different from the expected chance level (probability = 0.5). A decrease below chance level was only found for rotations at 135° and 180° in the palmar view, consistent with the predictions of the dual-process model (**Panel C**). **Panel C** shows the simulated predictions for the single- and dual-process models (see corresponding text for a detailed explanation on model parameterization.

### The ‘biomechanical constraints’ effect was present and more marked in the palmar view

This analysis included 62,121 trials. The GAMM identified a clear interaction between Direction and Hand View. In both views, the medial directions showed a general advantage over the lateral directions, consistent with a ‘biomechanical constraints’ effect. However, the effect was larger in the palmar view. For the palmar view, there was a consistent advantage of medial directions over lateral directions from 295ms onwards, reaching its peak at 462ms (difference = 0.18 [0.11, 0.25], p_FDR_ = 2.89×10^-6^), slowly decreasing afterwards, but never overlapping 0 (Fig. 5). In contrast, for the dorsal view, the difference between medial and lateral directions was found in the period 349-918ms (although it did not hold after correcting for multiple comparisons, p_FDR_ > 0.05), reaching its peak at 446ms (difference = 0.09 [0.02, 0.16), p_FDR_ = 0.09), linearly decreasing afterwards, the 95%CI gently overlapping 0 (Fig. 5).

**Figure 5.**
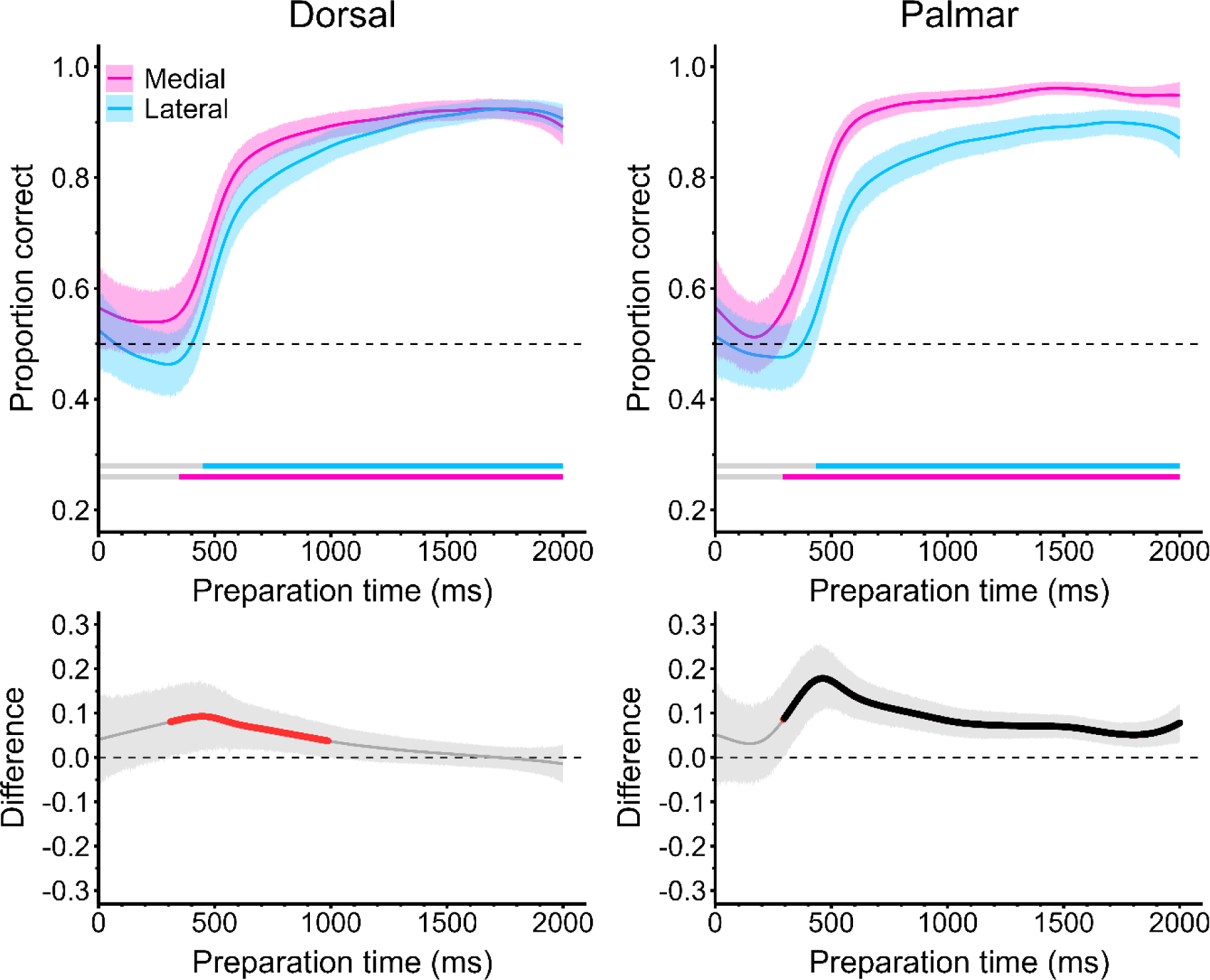
Information processing of medial and lateral directions, illustrating the ‘biomechanical constraints’ effect, for the dorsal view (left panel) and the palmar view (right panel). In the upper plots, chance level (probability = 0.5) is shown by the dashed line. Horizontal bars show when the condition is different from the expected chance level. Underneath each plot, the difference in the probability of being correct between medial and lateral is plotted across Preparation Time, positive values indicating differences in favour of medial directions, and negative in favour of lateral directions. The highlighted red and black regions in the lower panels show where the difference was p < 0.05 and p_FDR_ < 0.05, respectively.

## DISCUSSION

Traditional reaction time paradigms face limitations as they provide only data from the endpoint of information processing, which can be difficult to interpret due to inherent speed-accuracy trade-offs. We overcame those limitations in the HLJT by employing a ‘forced response’ paradigm, which allowed us to study the evolution of the time-course of stimulus processing with greater precision. Unsurprisingly, the results showed a strong effect of stimulus rotation, illustrating that the absolute rotation angle increased the time needed to process the stimulus. However, we found that this effect was non-monotonic, with the largest difference appearing when switching from 90° to 135°. Importantly, the effect of rotation was conditional on the view of the hand, which illustrated that palmar and dorsal views were indeed processed in fundamentally distinct ways for more extreme rotations. Finally, we also identified the presence of the ‘biomechanical constraints’ effect, which was primarily driven by the palmar view, in line with the idea that there are qualitative differences in the processing of palmar and dorsal views.

### The time-course of information processing

One of the strengths of our study is that we were able to determine *when* the differences between conditions are more marked over the time-course of processing, at the behavioural level. As, to the best of our knowledge, our study is the first to examine this time-course at the behavioural level in the HLJT, we discuss our findings in relation to studies using neurophysiological approaches, which have provided insights into the processing time-course in this task.

In the case of stimulus rotation, our results are broadly in agreement with the previous neurophysiological literature. We observed that the peak of the differences was consistent at around 450-500ms for all pairwise comparisons. This is consistent with data from electroencephalography (EEG) regarding the “Rotation-Related Negativity” (RRN) phenomenon in mental rotation tasks (Heil, 2002; Peronnet & Farah, 1989). The RRN is a parietal response where the amplitude of the P300 component negatively deflects as a function of the amount of rotation required (Ben-Shachar & Berger, 2024), and is present in the HLJT (Osuagwu & Vuckovic, 2014; ter Horst et al., 2012; Yu et al., 2020). Our results are also in agreement with subthreshold Transcranial Magnetic Stimulation studies showing that changes in reaction times in the HLJT occur when the stimulation is applied over the motor cortex between 100-500ms after stimulus onset (Pelgrims et al., 2009, 2011). While recent studies propose that mental manipulation of the stimulus occurs at a later stage (around 750ms; Davis et al., 2024), stimulation over the parietal lobe increased reaction times when applied as early as 250ms. This agrees with the time-course of stimulus processing we observed, as for stimuli with minimal rotations (0° and 45°), the approximate time to raise above chance level was around 300-350ms. This is again consistent with EEG literature showing that mental rotation of body parts or whole-body figures occurs around 310-380ms (Overney et al., 2005), although it has to be taken into account that ‘forced response’ paradigms often show reaction times 80-100ms faster than traditional ‘reaction time’ paradigms (Haith et al., 2016; Hardwick et al., 2022).

### The non-monotonic effect of stimulus rotation

Consistent with previous studies, our results indicate that the time required for stimulus processing generally increases with the absolute rotation applied. However, a non-monotonic effect was observed. This is consistent with previous studies that have shown that angular disparity of *two hands* on screen is proportional to reaction time (i.e. the slope is close to 1), but stimulus rotation of only *one hand* is not (Hoyek et al., 2014; Mibu et al., 2020). When one hand is presented, non-linear relationships between reaction time and stimulus rotation can occur (Bek et al., 2022; Conson et al., 2020; Ionta et al., 2007). For example, it is common to observe very little or no increases in reaction time from the “fingers upward” orientation (0°) to the next possible rotation (usually 30° or 45°), whereas larger increases are typically observed the closer the rotation gets to the “fingers downward” or maximum rotation (180°). A similar pattern was observed in the present study, with no discernible differences between 0° and 45°, but moderate-to-large differences for the rest of comparisons. Overall this evidence contrasts with traditional paradigms in mental rotation (which use two stimuli on screen), as these have consistently demonstrated that the increase in reaction time is relatively proportional to the angular disparity of stimuli (i.e., there is a *linear* relationship between the two) (Harris et al., 2000; Heil & Rolke, 2002). However, judging the similarity of two rotated stimuli (regardless of whether they are hands or objects) and judging the *type* of one rotated stimulus (e.g. laterality for hands, canonical vs. mirror-reversed forms for letters) are not technically the same task. Therefore, the expected functional relationship between reaction time and stimulus rotation could be different in these two scenarios, partially because the latter requires stimulus recognition whereas the former does not (Searle & Hamm, 2017).

Although our results are consistent with a non-monotonic relationship, the largest differences were observed when shifting from 90° to 135°, while further rotation (from 135° to 180°) had a relatively smaller effect. We interpret this under the ‘allocentric vs egocentric’ framework (see next section for details), which suggests a point exists where the processing of hand stimuli transfers from an egocentric to an allocentric reference frame. This is consistent with the proposal that there is a critical inflexion point at an angle between 90° and 135° which qualitatively changes how the stimuli are processed, from an egocentric to an allocentric frame of reference (Brady et al., 2011; Shmuelof & Zohary, 2008). In fact, a parallel body of evidence provides a similar framework, framing this shift in terms of the distinction of interpreting the stimulus as ‘self vs others’ (Ferri et al., 2011). In this model, egocentric processing (i.e., processing the hand as belonging to the individual’s own body) would lead to a ‘self-advantage’ compared to allocentric processing (i.e., processing the hand as belonging to another individual’s body) (Frassinetti et al., 2009). We note that both frameworks are largely equivalent in their predictions for this task.

All the above would be partially against the classical assumption in mental rotation that the stimulus is always rotated through the shortest angle to the upright (Searle & Hamm, 2017). This illustrates the typical, relatively symmetrical ‘V’ shape of reaction times plotted against angular disparity in mental rotation. An alternative explanation could relate to the fact that ‘same-different’ tasks (two stimuli) mainly involve visuo-spatial transformations, therefore relying primarily on visual processing, while ‘laterality judgement’ tasks of hands (one stimulus) could involve both visual *and* motor processing (or a mixture between the two) (Mibu et al., 2020). Nonetheless, we note there is also evidence that motor processes contribute to mental rotation in general (Wexler et al., 1998).

### Differences in processing for palmar and dorsal views

An important finding of our study was a strong interaction between stimulus rotation and hand view, illustrating a fundamental difference in the processing of palmar and dorsal views at more extreme rotations. Moreover, our data are in support of a strong interaction between the magnitude of the ‘biomechanical constraints’ effect and the view of the hand that is presented, showing that the effect is much stronger for palmar compared to dorsal views.

Two relatively overlapping frameworks could explain those findings. Firstly, it would be more natural for dorsal views to trigger processing using a ‘visual scanning’ approach, based on asymmetries in the position of the thumb (Conson et al., 2021). In this ‘visual vs motor’ framework, cognitive processes would be different between dorsal and palmar views primarily because of their low-level properties. Supporting evidence comes from studies showing that when the position of the thumb is manipulated so that asymmetry is removed in a dorsal view, information processing drastically changes from visual to motor processing (Conson et al., 2021; Hoyek et al., 2014). The shift in information processing is critical, to the extent that the ‘biomechanical constraints’ effect (which here is considered a hallmark of motor processing) critically emerges in dorsal views only when the thumb is ‘absent’ (Conson et al., 2021). In contrast, manipulation of the thumb position does not seem to affect the ‘biomechanical constraints’ in a palmar view, which would be *always* processed using a motor strategy. This suggests that thumb position ‘anchors’ processing in the dorsal view to a visual strategy and prevents other strategies (e.g., motor) from emerging. However, while this framework is useful to explain general differences between hand views, it does not offer a direct explanation about why hand views appear to be processed fundamentally differently only for the more extreme rotations.

On the other hand, the difference between palmar and dorsal views, generally *and* at extreme rotations, can be interpreted under the ‘allocentric vs egocentric’ framework (Brady et al., 2011; Wei-Dong et al., 2008) or the ‘self vs others’ framework (Ferri et al., 2011). These two models posit that there are hand postures that more naturally trigger processing from a first-person (egocentric) perspective, for example hands in the upward orientation (0°), whereas there are others that more automatically trigger third-person (allocentric) perspectives, for example the downward orientation (180°). Should this be the case, the key difference between the two hand views is the shift from egocentric (self) to allocentric (others) occurring only in the dorsal view at rotations greater than 90°. For such rotations, the most efficient and natural strategy would be the allocentric perspective (Nagashima et al., 2019; Shmuelof & Zohary, 2008). However, because palmar views are strongly associated with an egocentric perspective *regardless of stimulus rotation*, a penalty is paid for the most extreme angles in this view at the early stage of the processing time-course. This is illustrated by the decrease in accuracy below chance level observed in the present study around 300-500ms for 135° and 180° in the palmar view only. At that stage, rapidly-available spatial information may be prioritised over anatomical information, which requires more time to be processed (Waltzing et al., 2024). This creates a ‘conflict’ which is better captured by the dual-process model according to our computational modelling approach and reflected by an initial decrease in accuracy. However, this conflict is quickly overcome as anatomical information becomes available (from around 500ms onwards, see Figure S3D-E), which in turn makes the palmar view easier to process than the dorsal view, especially at 180°.

In summary, while our results are generally in line with the ‘visual vs motor’ framework, the ‘allocentric vs egocentric’ and ‘self vs others’ frameworks provide further plausibility about the differences in information processing between hand views as a function of stimulus rotation. Future studies could combine our paradigm with neurophysiological techniques to fully elucidate the neural underpinnings of these findings.

### The ‘biomechanical constraints’ effect

Results from our present manuscript demonstrated participants needed more time to accurately process stimuli with lateral compared to medial rotations, consistent with the presence of a ‘biomechanical constraints’ effect. The exact nature of the ‘biomechanical constraints’ effect is still debated, and it has been reviewed elsewhere (Moreno-Verdú & Hardwick, 2022). Briefly, while it has traditionally been considered to represent a hallmark of the use of motor imagery (Conson et al., 2020; Parsons, 1987), other work has proposed it could be attributed to implicit perceptual knowledge of anatomical constraints, rather than evidence that individuals are manipulating the mental representation of their *own* hand per se (Vannuscorps et al., 2012). Our results are in line with previous reports showing that the effect is stronger or only statistically significant in the palmar view (Conson et al., 2021; Mibu et al., 2020). Furthermore, our findings provide insight into the shape of the time-course of this effect, showing that the medial-to-lateral advantage is consistent across information processing, peaking at around 450ms. This is in line with the RRN phenomenon found in the HLJT. In fact, EEG data show that a key difference between medial and lateral directions was that the former produced significantly smaller RRN than the latter, providing neurophysiological evidence of the medial-to-lateral advantage (ter Horst et al., 2012). Moreover, other authors have found that manipulating the actual position of the individual’s hand to a position congruent with lateral rotations made them able to overcome the medial-to-lateral advantage. Interestingly, it led to a reduction in the RRN for lateral rotations (Jongsma et al., 2013), which the authors interpreted as evidence in favour of kinesthetic motor imagery being responsible for this effect (as it is the most plausible hypothesis given that visual processing would not be affected by actual body posture). Collectively, the cited literature agrees with our data, which showed a processing ‘benefit’ of medial directions over lateral directions (as illustrated by an earlier rise of the speed-accuracy trade-off function), but no difference in the actual shape of the information processing time-course. As such, we hope our results will stimulate further research into the neural and conceptual underpinnings of the ‘biomechanical constraints’ effect.

### Strengths and Limitations

This study has several limitations. ‘Forced response’ paradigms are useful to decompose the time-course of information processing, yet they also require multiple trials across a broad range of different time points. Therefore, a much larger number of trials is needed, making the paradigm more time-consuming and potentially fatiguing. Because of this, we focused on collecting enough trials per condition to allow analysis of our main questions-of-interest, while also ensuring the length of the overall experiment did not become too long, making it easier for participants to complete the task while minimising attrition and fatigue (particularly as we conducted the study fully remotely). In turn, this limited our ability to conduct some highly-specific analyses due to a lack of trials in that specific condition – for example, we could not test the effect of stimulus rotation on the ‘biomechanical constraints’ effect (i.e., whether the magnitude of the medial-to-lateral advantage varies depending on the specific rotation angle at which it is measured).

In addition, forced response protocols require participants to follow specific instructions in terms of response timing. This makes it necessary to provide timing feedback on a trial-by-trial basis, to ensure participants are following the instructions given. This in turn makes the provision of accuracy an important aspect of the task, as otherwise participants would only focus on producing well-timed (but not necessarily correct) responses. Since both aspects are important in forced response protocols, we included both types of feedback, which is a deviation from traditional HLJT paradigms (though we note there are situations where trial-to-trial feedback is provided in traditional reaction time based HLJT studies. Nonetheless, the fundamental effects of the task were replicated unambiguously in our protocol, which builds upon the previous literature consistently, showing robust behavioural phenomena even in the presence of small deviations like feedback provision.

Similarly, given that the HLJT only has two possible responses (left or right), and that all our participants were right-handed, a response preference for their dominant hand was found when they had no time to process the stimulus. This made the baseline level for left and right stimuli substantially different and therefore prevented us from further analysing the effect of laterality and its interaction with the other factors. However, give that our central question was to examine differences in information processing in the task overall, questions related to laterality/hand dominance were secondary to our investigation. Furthermore, our study is also valuable as a proof-of-concept that this type of behavioural paradigm can be feasibly conducted for the HLJT and fully remotely.

We also acknowledge that the choice of splitting participants in two groups based on hand view (which again was a decision made for feasibility purposes), might have influenced the results. However, our statistical analyses are robust to this fact, reducing the possibility of a between-group difference in socio-demographic characteristics or imagery ability being the source of the effects found. Splitting participants into two groups therefore provided a pragmatic solution to completing the study when taking into account the overall duration of the experiment, and the difficulties associated with completing a full crossover-counterbalanced design with a remote sample.

Finally, we did not include a control group or condition using a classic mental rotation task with non-biological stimuli (e.g. letters), as we were primarily interested in providing insight into how information is processed in the HLJT, and information processing in those tasks has been assessed elsewhere (Gardony et al., 2017; Jansen et al., 2020; Krause et al., 2021). However, we note that it is common practice for HLJT studies to consider only hand stimuli, as there is no clear ‘control’ stimulus which would match the main characteristics of a human hand.

## CONCLUSIONS

Information processing in the HLJT depends critically on stimulus rotation and the view of the hand that is presented. Greater stimulus rotation increased processing time, but this effect was not constant across the range of possible rotation angles. This non-linear effect should be considered when designing HLJT experiments. Furthermore, our results indicate a fundamental difference in processing of stimuli in palmar and dorsal views when presented at extreme rotations, illustrating a strong interaction between these two factors. This agrees with evidence suggesting that the two views may entail relatively separate cognitive strategies. The strength of the ‘biomechanical constraints’ effect substantially varies depending on the view of the hand, providing further evidence in this regard. Our results reveal the time-course of information processing during the HLJT in unprecedented detail, elucidating the cognitive processes involved in this task, and providing insights into the broader topic of motor imagery.

## ACKNOWLEDGEMENTS

This study is funded by the FNRS CR Fellowship (FNRS 1.B359.25) awarded to MMV. MMV, BMW, SMcA and RMH are supported by an FNRS MIS Grant (F.4523.23). EVC is supported by an FNRS ASP fellowship (1.AB19.24). GH is supported by an FNRS CDR award (J.0084.21).

## Notes

https://osf.io/z6b4d/

### Competing Interest Statement

The authors have declared no competing interest.

### Summary of Updates

A revision has been made to include literature in the Introduction and Discussion that was relevant but nor previously identified. In addition, changes have been made to the Introduction to highlight the novelty of the study. Finally, a power analysis has been conducted and reported in Supplementary Materials. Minor modifications have been made to Fig. 1 and throughout the text.

