## Supplementary Materials for "Information processing in the Hand Laterality Judgement Task: Fundamental differences between dorsal and palmar views revealed by a “Forced Response” paradigm"

### Details on Generalised Additive Mixed Models (GAMMs)

Additive models are a type of statistical model which allow to accommodate non-linear associations between predictors and response. Generalised Additive Mixed Models are an extension of simple additive models where the response variable is not continuous (e.g. binary), therefore known as "Generalised", and random effects are included in the model, therefore known as "Mixed". All additive models allow to flexibly make predictions about the dependent variable without assuming linear relationships with the predictors thanks to smoothing functions (also known as "splines"). Splines are actually constructed of many smaller functions (basis functions), and each smooth is the sum of a number of basis functions. Each basis function is multiplied by a coefficient, each of which is a parameter in the model. Therefore, a smooth can take virtually any possible shape, being a flexible way to capture non-linear associations. There are a number of types of splines each with its own properties (Wood, 2003).

However, additive models' higher flexibility can potentially present issues because they can overfit the data and become too 'wiggly' (Pedersen et al., 2019). There are several ways to control the 'wiggliness' in additive models. One of them is to penalise the change in the slope (i.e. a penalty to the second derivative or second-order penalty). This is done to avoid overfitting and can be performed controlling the number of basis functions that make up the smooth function, which corresponds to k parameter when working with the 'mgcv' package in R.

In this study, low-rank thin plate regression splines were used (Wood, 2003). While building the models, we also considered cubic regression splines but fits between these and thin plate splines were similar, so the latter were selected as they are more flexible. To prevent overfitting, the 'wiggliness' of the GAMMs was controlled by a second-order penalty via iteratively selecting the appropriate value of the k parameter. Model fit was first assessed based on its effective degrees of freedom (edf), k-index, and the corresponding p-value; if a problem was detected (edf approached k, k-index < .95 and $p < 0.05$), a model with k+5 was built (Wood et al., 2016). The process was repeated iteratively until the above-mentioned indexes were observed. Goodness-of-fit across all candidate models was performed by Akaike's Information Criterion (AIC), and the model with the smallest AIC was finally selected. For a given selected model, model misspecification was first screened by inspecting residual plots, zero-inflation and outliers using the 'DHARMa' package.

### Methodological checks

The groups were not different in the percentage of available trials for analysis (Palmar = 97.5 ± 1.56%; Dorsal = 97.02 ± 1.81%; $t_{(52)} = 1.03$, $p = 0.309$; Fig. S1A). There was a lower number of available trials for actual Preparation Times >2000ms in both groups (Fig. S1B). Therefore, subsequent predictions were limited to the range 1-2000ms, which was in line with the experimental paradigm (20 to 1980ms). Participants generally responded according to the given instructions, and in most trials the response time was within the ± 100ms limit indicated (Fig. S1C). However, there were also many trials beyond this limit, which highlighted the need to analyse all available trials considering the Preparation Time participants actually used to process the stimuli, regardless of the theoretical Preparation Time they should have employed.

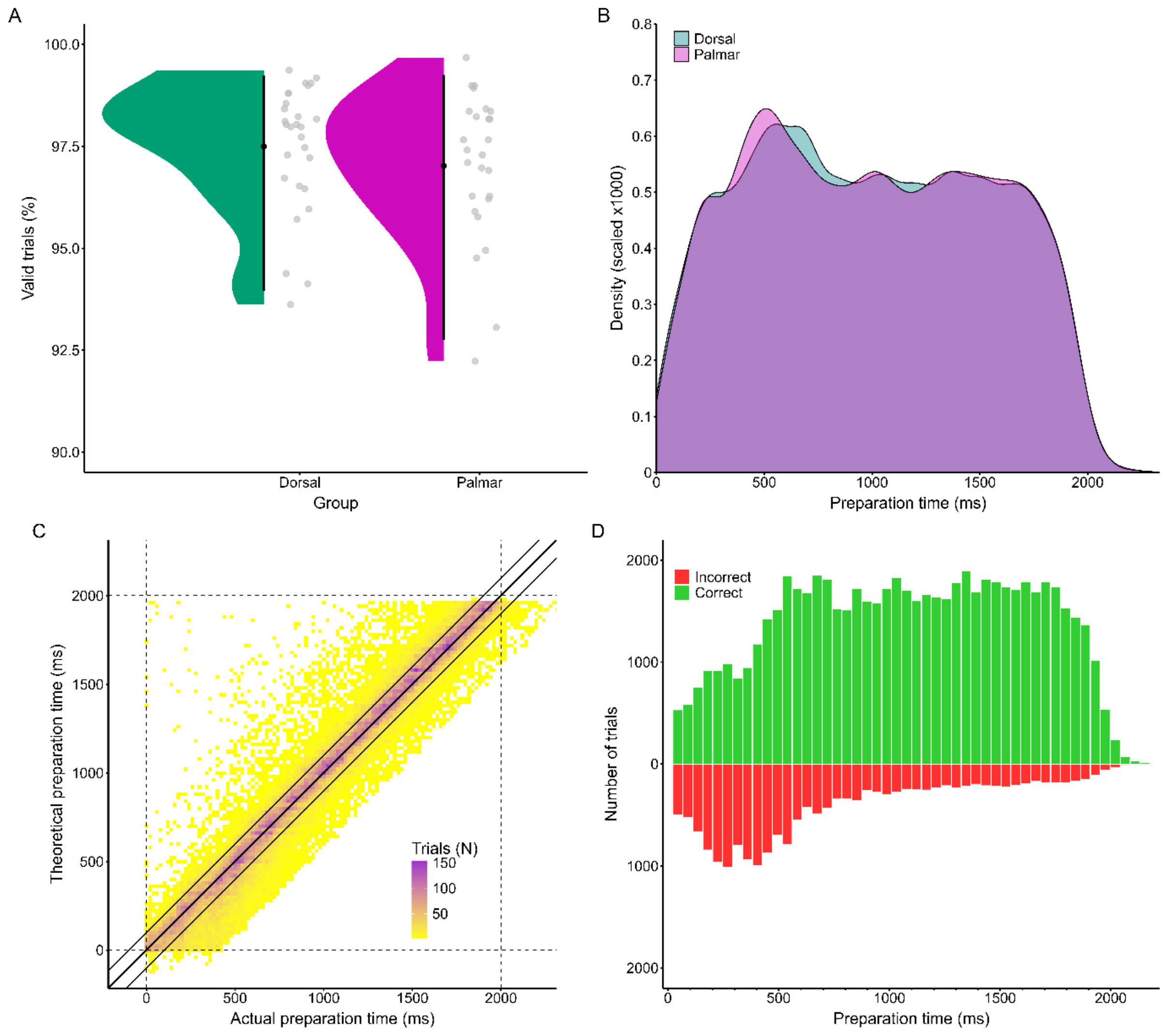


**Supplementary Fig. 1. Panel A** shows the percentage of valid trials analysed by experimental group. **Panel B** shows a kernel density plot representing the number of trials available for analysis according to the Preparation Time and Hand View. **Panel C** is a heatmap which shows the relationship between the actual Preparation Time and the intended (theoretical) Preparation Time. If participants had perfectly synchronised their response exactly at the same time as the last tone, all points would lie along the diagonal 'equivalence' line (solid black line). In reality, timing was considered accurate as long as it was within +-100ms of the intended preparation time (represented by the parallel lines either side of the equivalence line). The heatmap therefore illustrates that participant responses generally respected this timing, but that there was variability that should be taken into account. **Panel D** shows the total distribution of correct and incorrect trials throughout Preparation Time. At very early processing, the distributions are approximately the same (i.e. accuracy = 0.5), but as more processing time is available, the number of incorrect responses decreases and the number of correct responses increases.

### Pairwise comparisons between Rotation Angles for each Hand View separately

This analysis compared the speed-accuracy trade-off for each pair of sequential Rotation Angles (e.g. 0° and 45°, 45° and 90°, etc) for the dorsal and palmar view separately. The analysis was based on predictions from the same GAMM.

Differences between 0° and 45° were not found in the dorsal view (difference < 0.01, $p_{FDR} > 0.99$; Fig. S2A), or the palmar view (difference < 0.02 , $p_{FDR} > 0.56$; Fig. S2E) at any time period. Differences between 45° and 90° in the dorsal view were found from 423ms onwards, reaching the peak at 457ms (difference = 0.08 [0.01, 0.16], $p_{FDR} = 0.03$) and decreasing afterwards, approaching 0 but the 95%CI never overlapping (Fig. S2B). In the palmar view, differences between 45° and 90° were found in the period 313-1664ms, reaching the peak at 406ms (difference = 0.15 [0.08, 0.23], $p_{FDR} = 0.0004$) and decreasing afterwards, approaching 0 but the 95%CI never overlapping (Fig. S2F). For 90° and 135°, in the dorsal view differences were observed in the period 448-1921ms, the peak being at 591ms (difference = 0.18 [0.11, 0.25], $p_{FDR} = 3.96x10^{-6}$), decreasing afterwards, approaching 0 (Fig. S2C). For the palmar view, differences between 90° and 135° were found between 324-1663ms and 1718ms onwards, reaching the peak at 450ms (difference = 0.28 [0.19, 0.36], $p_{FDR} = 1.6x10^{-9}$; Fig. S2G). Finally, differences between 135° and 180° were found from 532ms onwards in the dorsal view, reaching the peak at 746ms (difference = 0.17 [0.08, 0.26], $p_{FDR} = 0.0007$), and maintaining afterwards, never approaching 0 (Fig. S2D). However, in the palmar view, differences were found in two clusters (493-678ms, and 1410-1679ms), with the peak found at 557ms (difference = 0.12 [0.03, 0.22], $p_{FDR} = 0.13$; Fig. S2H), but none of these comparisons survived after correcting for multiple comparisons ($p_{FDR} > 0.05$).

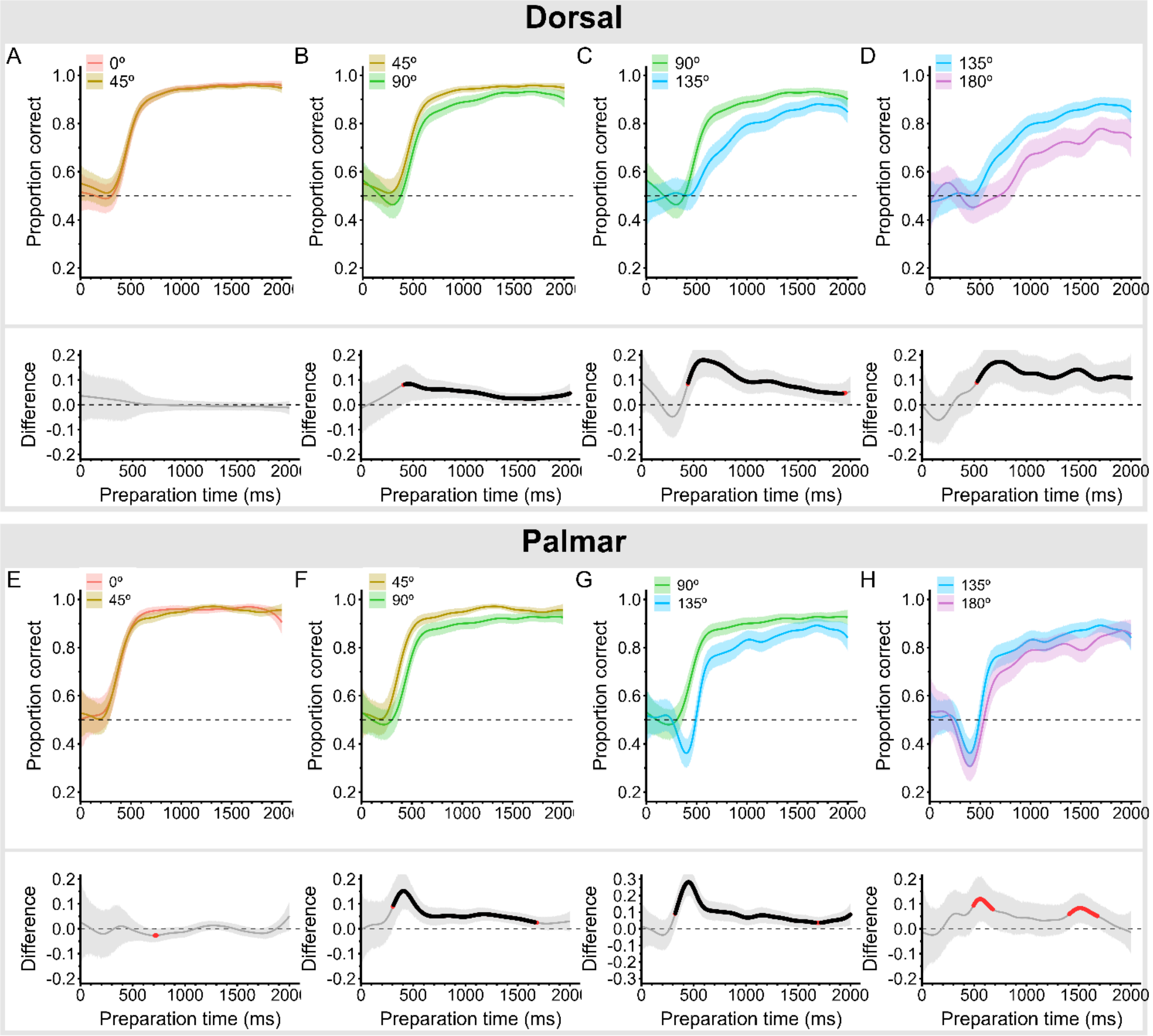


**Supplementary Fig. 2.** Speed-accuracy trade-off plots for each Rotation Angle pairs depending on Hand View (Panels A-D for the dorsal view and Panel E-H for the palmar view). Underneath each panel, the difference between each pair is plotted as Difference = Angle 1 – Angle 2, and therefore positive differences illustrate an advantage of the less rotated stimulus. The highlighted red and black regions in the lower panels show where the difference was $p < 0.05$ and $p_{FDR} < 0.05$, respectively.

### Comparison between Hand Views for each Rotation Angle separately

This analysis compared the speed-accuracy trade-off for each Hand view at each Rotation Angle separately. The analysis was based on predictions from the same GAMM.

For a rotation of 0° (Fig. S3A), the palmar view showed an advantage over the dorsal view in the period 326-820ms, with a peak at 415ms (difference = 0.15 [0.08, 0.23], $p_{FDR} = 0.0006$). For a rotation of 45° (Fig. S3B), the palmar view showed an advantage in the period 345-590ms, with a peak at 413ms (difference = 0.15 [0.08, 0.23], $p_{FDR} = 0.0006$). For a rotation of 90° (Fig. S3C), the palmar view showed an advantage in the period 412-612ms, with a peak at 454ms (difference = 0.09 [0.007, 0.16], $p_{FDR} = 0.44$), but the difference did not survive after correcting for multiple comparisons ($p_{FDR} > 0.05$). For a rotation of 135° (Fig. S3D), the dorsal view showed an advantage in the period 324-460ms, with a peak at 396ms (difference = -0.14 [-0.22, -0.06], $p = 0.09$), whereas the palmar view showed an advantage in the period 563-802, with a peak at 624ms (difference = 0.11 [0.03, 0.19], $p_{FDR} = 0.09$), but neither survived after correcting for multiple comparisons ($p_{FDR} > 0.05$). Finally, for a rotation of 180° (Fig. S3E), the dorsal view showed an advantage in the period 286-466ms, with a peak at 383ms (difference = -0.15 [-0.24, -0.06], $p_{FDR} = 0.009$), whereas the palmar view showed an advantage between 573-1469ms and 1689ms onwards, with a peak at 715ms (difference = 0.19 [0.11, 0.28], $p_{FDR} = 0.001$). Importantly, the difference in favour of the dorsal view at 180° was due to the dip in accuracy of the palmar view instead of a raise from chance level in the dorsal view. Conversely, the difference in favour of the palmar view was due to an advantage of this view.

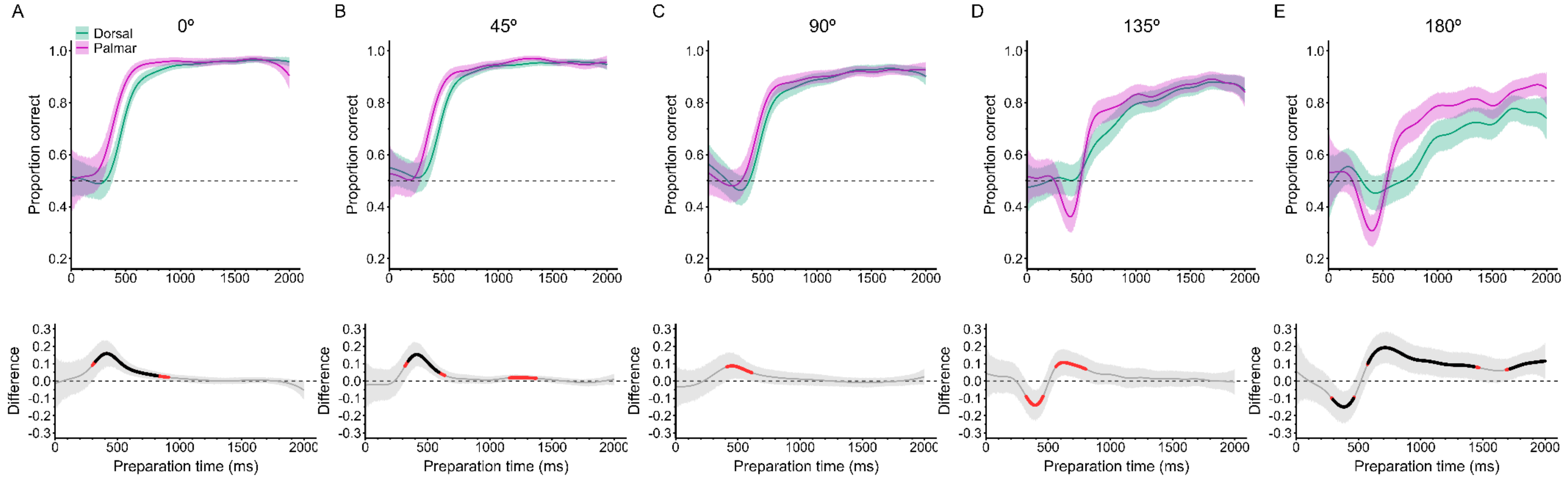


**Supplementary Fig. 3.** Speed-accuracy trade-off plots for each Hand View pair depending on Rotation Angle. Underneath each panel, the difference is plotted, positive values indicating an advantage of the palmar view, and negative values an advantage of the dorsal view. The highlighted red and black regions in the lower panels show where the difference was $p < 0.05$ and $p_{FDR} < 0.05$, respectively.

### Handedness produced a 'preference' for making 'random responses' with the right hand

The forced response paradigm assumes that if participants do not have enough time to process a stimulus, they should respond at random (i.e. in the present experiment, a 50-50 chance of responding with the left or right hand). A separate GAMM was intended to be run considering the effect of laterality (i.e. analysing right and left stimuli independently or comparing the effect of laterality on the other relevant factors in the model). However, while running the analysis we identified different baseline levels for chance in the short Preparation Time period (0-400ms) between right and left stimuli. Right stimuli were consistently above chance level, whereas left stimuli were consistently below chance level, across all conditions (see for example Fig. S4 from sample data).

As participants did not have enough time to process the stimulus in this period, we further inspected whether this difference was due to a 'response preference', i.e. when responding 'at random', participants generally preferred to respond with their dominant (right) hand. Such an effect would be analogous to 'finger effects' sometimes present in the broader literature on reaction time. While participants were not generally more inclined to respond with their dominant (right) hand (Fig. S5A showed that overall, the proportion of right and left responses was at 0.5), further analysis indicated that this varied across the processing time-course. As expected, participants showed the preference in both the palmar and dorsal groups only at the shortest Preparation Times (Fig. S5B), i.e., when they were spontaneously responding 'at random'. Importantly, we confirmed that this preference did not interact with actual accuracy by examining the proportion of correct responses for right and left stimuli across Preparation Time, which due to the design of our experiment should always been around 0.5 (Fig. S5C shows this was largely the case). Finally, we also ruled out the possibility that the preference was due to having a higher number of available trials responded with the right hand (i.e. due to rejecting more left-hand trials than right hand trials), which was not the case (Fig S5D).

Based on the above data, we considered it was not appropriate to include the effect of laterality in the main analysis, as the results could have been conditioned by the identified 'preference'. Therefore, the main results were presented collapsing across right and left stimuli, cancelling out the effect of laterality.

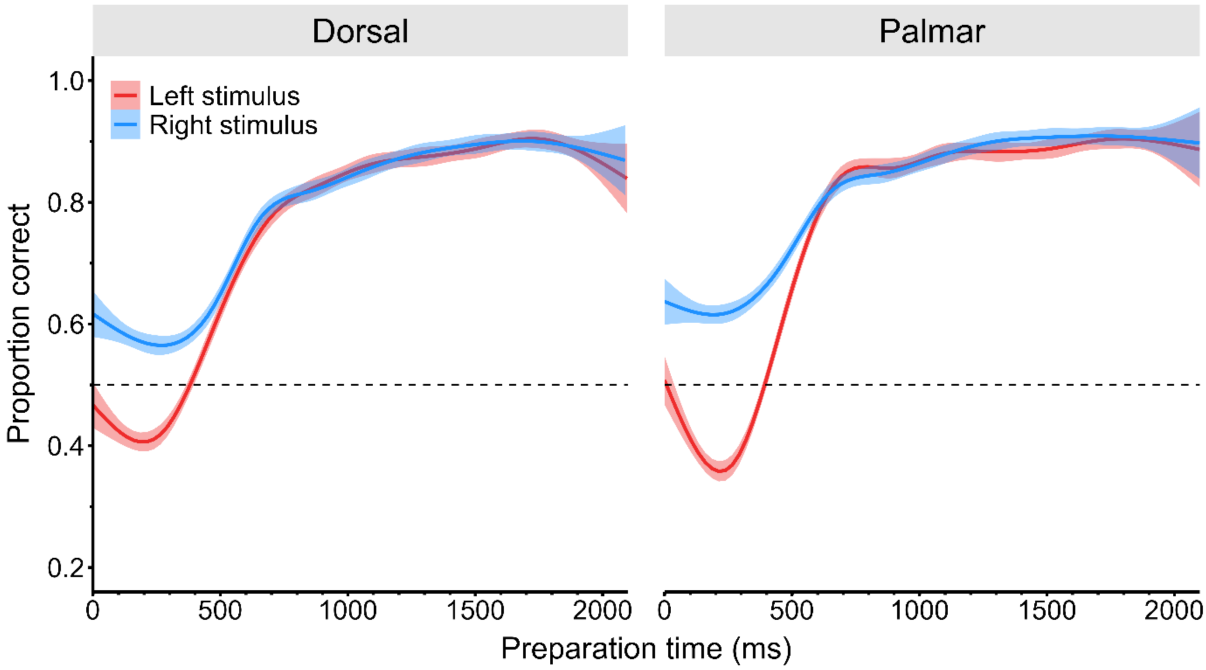


**Supplementary Figure 4.** Speed-accuracy trade-off plot splitting by Hand View and Stimulus Laterality. The chance level of right stimuli is above chance (dashed line: 0.5) across views in the early phase of the processing time-course, and the chance level for left stimuli is below chance level. This plot shows sample data, and no statistical modelling was applied.

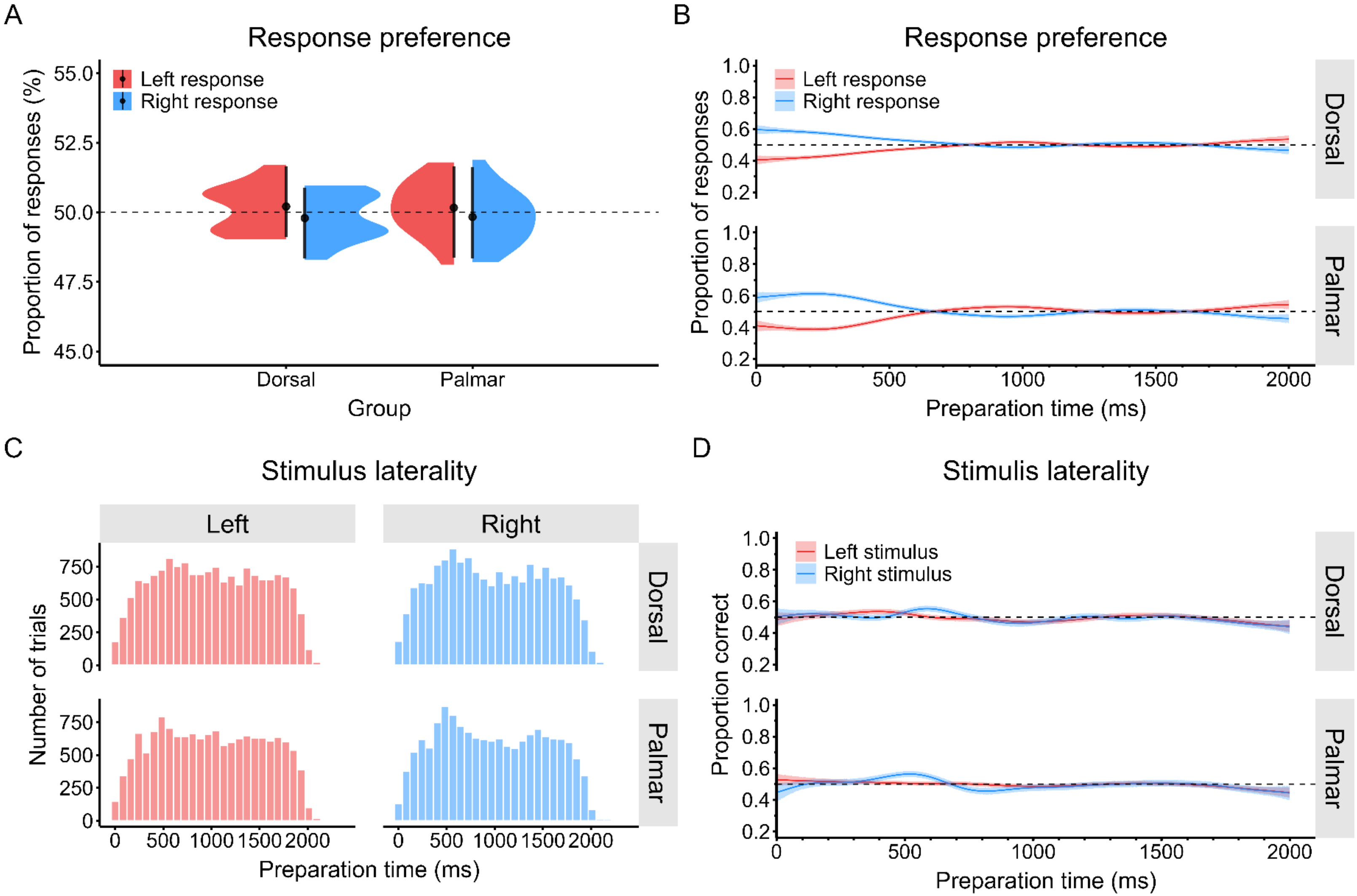


**Supplementary Figure 5. Panel A** shows the average proportion of actual responses throughout the experiment, for the right/left response in the palmar and dorsal groups. **Panel B** shows the actual response selection as a measure of the time available to prepare the response, illustrating the response 'preference' participants exhibited, as they used their right (dominant) hand to respond in the random period, when they did not have time to process the stimulus. **Panel C** shows the number of trials available for analysis for each level of Stimulus Laterality and Hand View, illustrating a similar distribution across conditions. **Panel D** shows the proportion of correct responses across Preparation Time. As expected, this proportion was approximately at chance level (probability = 0.5) throughout the range of Preparation Time.

### Power Analysis

We performed a simulation-based power analysis focusing on the key interaction between Rotation Angle and Hand View. We simulated, through binomial distributions, speed-accuracy trade-offs (SAT) considering the proportion of correct responses as the dependent variable and preparation time as the main continuous covariate driving information processing. As a fundamental effect, we simulated how the SAT varied as a function of Rotation Angle, whereby increasing stimulus rotation produced a large effect (i.e., Cohen's d = 0.8) on the slope of the SAT. For simplicity, we assumed this effect was monotonic across the angles of rotation. Based on the results of the present study, the main parameter that was affected by rotation angle was the centre of the slope (i.e., the midpoint between the two asymptotes in the SAT). Increasing stimulus rotation was assumed to shift the slope's centre, illustrating slower information processing as a function of rotation. To inform this amount, we extracted the data from our previous study which used identical stimuli and the closest experimental conditions, in a traditional reaction time "free" paradigm (Moreno-Verdú et al. 2025, doi: 10.1016/j.neuroscience.2025.02.056; data can be freely accessed at https://osf.io/8h7ec/). In that study the average difference between the consecutive angles (e.g., 0° vs 45°, 45°vs 90°, etc.) was 135.68 ms across Hand Views (101.87 ms for palmar and 175.15 ms for dorsal). Hence the amount of the centre's shift was obtained by multiplying the observed mean difference by the effect size, as a conservative expectation (135.68 * 0.8 = 108 ms for every 45° absolute increase, starting at 0° and finishing at 180° in steps of 45°). Two other parameters were affected by stimulus rotation, namely the final asymptote of the SAT and the steepness of the slope. The influence of Rotation Angle on these parameters was weaker than the centre of the slope (as can be seen in Fig. 1C of the present study). The average final asymptote (proportion correct = ~0.95) was expected to occur at ~1000ms and to slightly decrease with rotation angle, with an average decrease of ~2% (hence for 0° = 0.95, for 45° = 0.93, 90° = 0.91, etc.). The steepness of the slope was assumed to increase with the rotation angle, with a rate of ~2/ms for every increase in 45° of rotation. This means that for every 45° increment in angle, the rate at which accuracy improves with preparation time becomes steeper, reflecting a ~90% increase in the steepness per 45° of rotation. We note that these parameters were chosen to build SATs that were globally compatible with the ones obtained in the present study, but the parameters can be adjusted for other scenarios their specific values do not affect the main power calculation.

As the main power calculation, the effect of angle was assumed to be different between Hand Views (dorsal and palmar). To estimate the expected effect size, we conducted a frequentist re-analysis of our previous study (Moreno-Verdú et al. 2025, as above) focusing on the Angle x View interaction (both were within-participant factors in that study). We considered Reaction Time as the dependent variable in a linear mixed-effects model with random intercept for participant (mimicking our GAMMs); formula: Reaction Time ~ Angle x View x Laterality x Group + (1|ID). Note this model included Laterality (left/right) and Group (in-person/online) which are not of interest for the present analysis but were factors in the original study that we kept for consistency with the original analysis. The results indicated a statistically significant Angle x View interaction with a moderate-to-large effect size according to established benchmarks (Richardson 2011; Cohen 1969): $F_{(7,1177)} = 20.87$, $p = 1.3 \times 10^{-26}$; $\eta_p^2 = 0.11$, 95%CI [0.08, 0.14]; partial Cohen's f = 0.35 [0.29, 0.40]. Because this comparison was within-participants, but in the present study the comparison was between-participants, we calculated power for effect sizes ranging from moderate to large based on Cohen's conventions. For computational efficiency, calculations were obtained only for d = 0.5, d = 0.65 and d = 0.8 (d = 0.65 as the intermediate point between 0.5 and 0.8).

The main power calculation included simulating 1,000 datasets with varying sample sizes (from N = 30, in steps of N = 4, up to N = 102 participants total, half per view). We did that for each effect size, assuming each participant contributed with 1584 trials, as per our experimental design. For each participant independently, 99 preparation times were randomly sampled in steps of 20ms from a distribution ranging from 1 to 1980ms, with 8 rotational angles sampled twice each (right and left stimuli) per preparation time per participant. Therefore, 3 effect sizes x 19 sample sizes x 1,000 datasets were simulated overall. An example dataset is shown in Fig. S6.

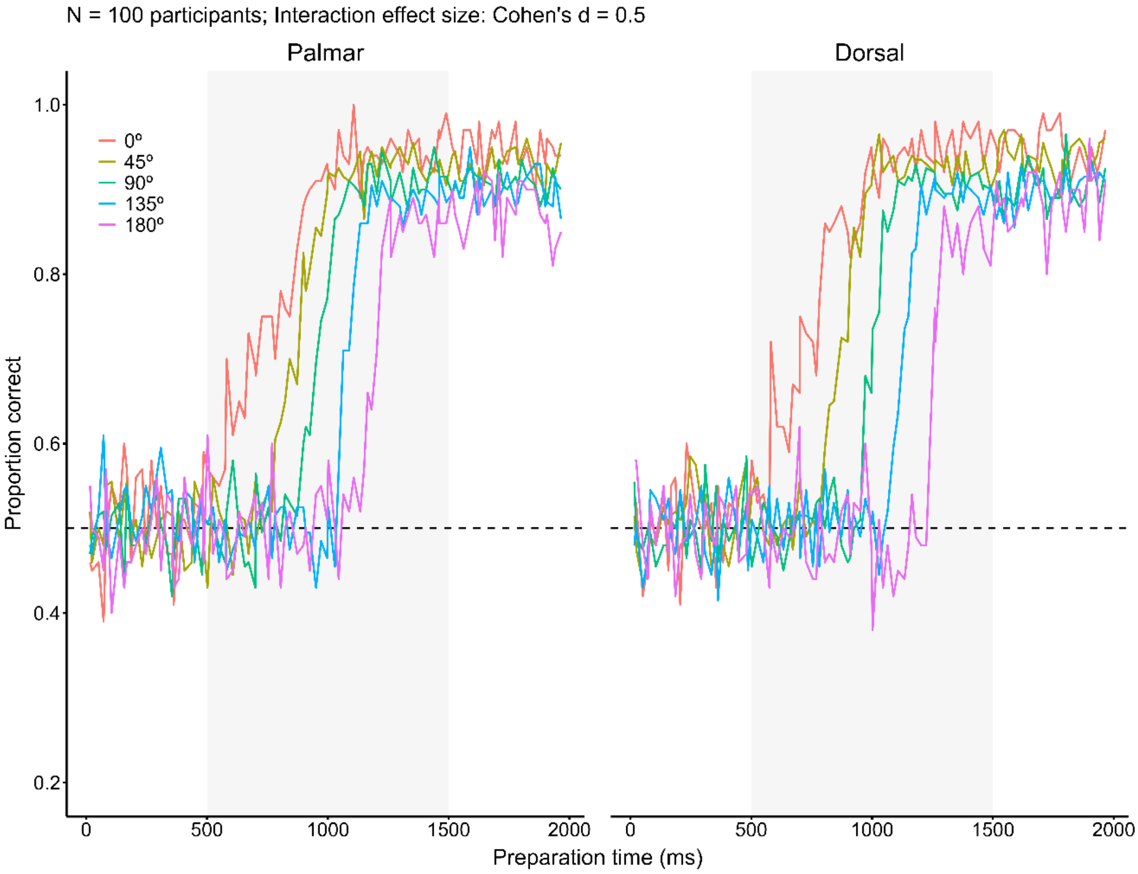


**Supplementary Figure 6.** Example dataset with N=100 total participants (half per view), and a moderate interaction effect size (Cohen's d = 0.5). The shaded grey area shows where the principal effect of the interaction can be seen. For the dorsal condition, the slope of the speed-accuracy trade-off is shifted towards the right as a function of rotation angle to a greater extent than for the palmar condition (i.e. slower information processing overall). The steepness of the slope and the final asymptote are also slightly affected by the interaction.

In GAMMs, each combination of levels for each factor in the interaction is given a separate smooth (i.e., a separate SAT in our scenario). Therefore, there is no direct way to derive the effect of interactions as with traditional ANOVA's omnibus tests. Because of that, the effect of the interaction was calculated using model comparison. For each complete dataset, two GAMMs were fitted, always with a random intercept for participant (irrelevant for the simulation as no random effects structure was included, but to keep consistency with the main analysis of the present study) and using a binomial distribution with a logit link function. The first model included smooth terms for the main effects of Rotation Angle and Hand View, but without their interaction. The second model included smooth terms for both main effects and their interaction. The models were fitted varying the number of basis functions to control data overfitting, via varying the k parameter (k = 20 as the main analysis, and k +/- 5 as a sensitivity analysis). For each dataset, the models were compared through a $\chi 2$ test, where a p-value < 0.05 (alpha = 5%) was considered evidence in favour of the model including the interaction. We derived power as the proportion of datasets that yielded a p-value < 0.05 for each sample size and effect size.

The results are shown in Fig. S7. To achieve the traditional nominal power of ~80%, N=38, N=54 and N=78 participants would be required for large, moderate-to-large and moderate effect sizes, respectively. To achieve a more conservative power of ~95%, N=46, N=66 and N>102 participants would be required for the corresponding effect sizes.

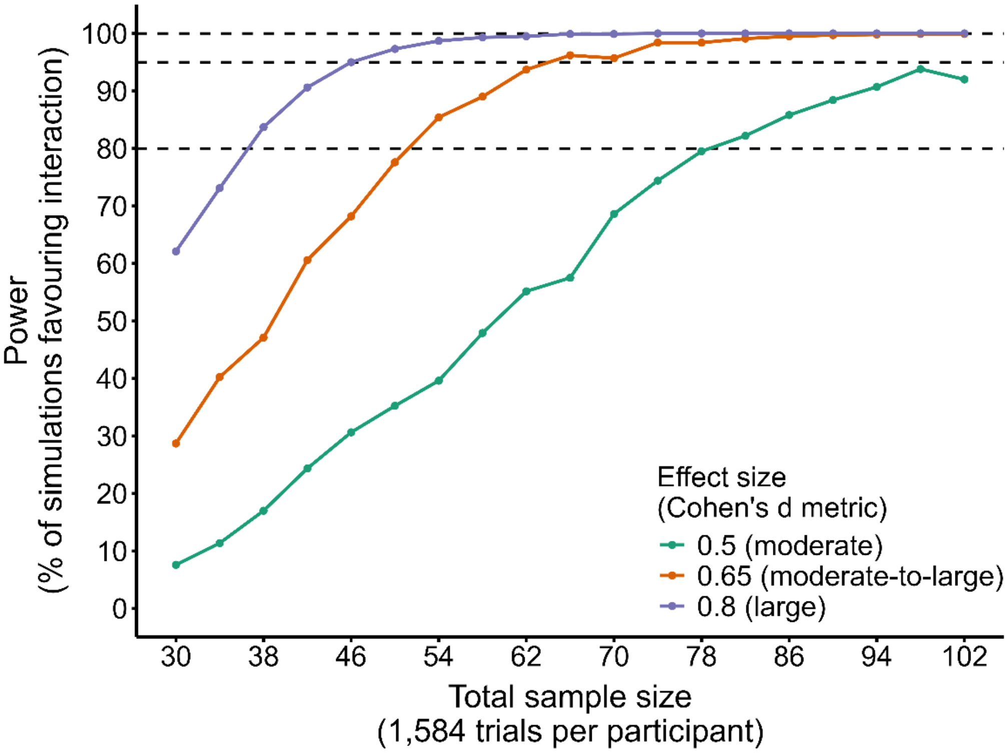


**Supplementary Figure 7.** Results of power analysis for the Rotation Angle x View interaction, for each effect size and sample size. Power is shown as the proportion of simulations yielding p-value < 0.05 in favour of the GAMM model with interaction, compared to the GAMM model with main effects but no interaction. GAMMs are fitted with k = 20.

The results were largely consistent across GAMMs with different k parameters (Fig. S8).

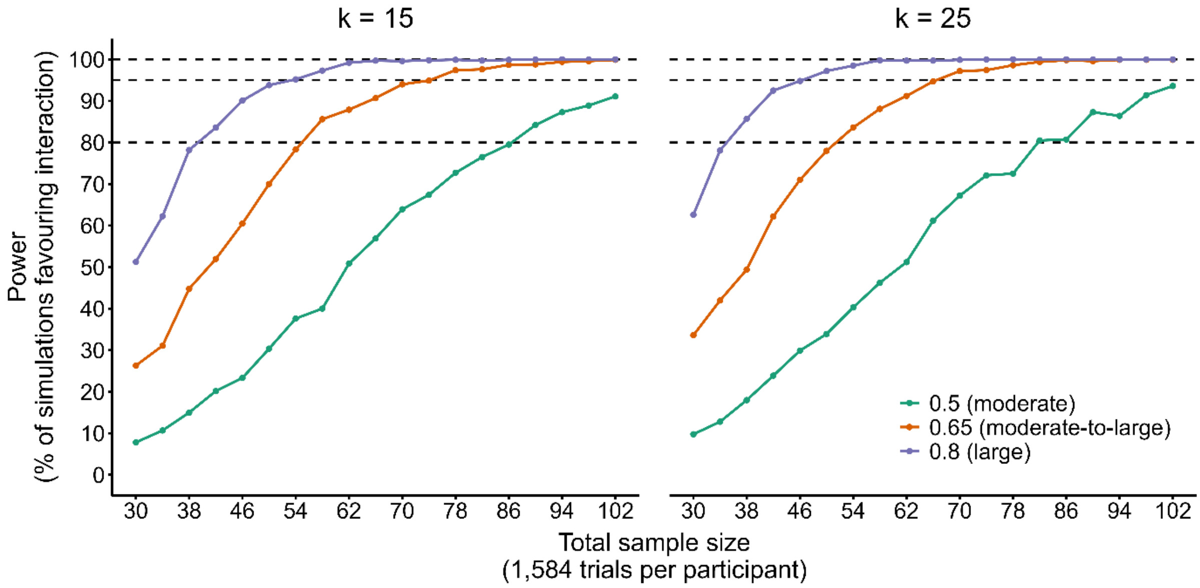


**Supplementary Figure 8.** Results of sensitivity analysis, varying the k parameter in the GAMMS.
